# Phylogenetic relationships among the clownfish-hosting sea anemones

**DOI:** 10.1101/560045

**Authors:** Benjamin M. Titus, Charlotte Benedict, Robert Laroche, Luciana C. Gusmão, Vanessa Van Deusen, Tommaso Chiodo, Christopher P. Meyer, Michael L. Berumen, Aaron Bartholomew, Kensuke Yanagi, James D. Reimer, Takuma Fujii, Marymegan Daly, Estefanía Rodríguez

**Affiliations:** Division of Invertebrate Zoology, American Museum of Natural History, New York, NY, USA; Sackler Institute for Comparative Genomics, American Museum of Natural History, New York, NY, USA; Department of Biological Sciences, Auburn University, Auburn, AL, USA; Department of Biology, University of Houston, Houston, TX, USA; Department of Invertebrate Zoology, National Museum of Natural History, Smithsonian Institution, Washington DC, USA; Division of Biological and Environmental Science and Engineering, Red Sea Research Center, King Abdullah University of Science and Technology, Thuwal, Saudi Arabia; Gulf Environments Research Institute, American University of Sharjah, Sharjah, United Arab Emirates; Coastal Branch of Natural History Museum and Institute, Chiba, Kastsuura, Chiba, Japan; Molecular Invertebrate Systematics and Ecology Laboratory, Department of Biology, Chemistry, and Marine Sciences, Faculty of Science, University of the Ryukyus, Nishihara, Okinawa, Japan; Tropical Biosphere Research Center, University of the Ryukyus, Nishihara, Okinawa, Japan; International Center for Island Studies, Kagoshima University, Kagoshima, Japan; Department of Evolution, Ecology, and Organismal Biology, The Ohio State University, Columbus, OH, USA

**Keywords:** Actiniaria, clownfish, symbiosis, mutualism, Actinioidea, Stichodactylidae

## Abstract

The clownfish-sea anemone symbiosis has been a model system for understanding fundamental evolutionary and ecological processes. However, our evolutionary understanding of this symbiosis comes entirely from studies of clownfishes. A holistic understanding of a model mutualism requires systematic, biogeographic, and phylogenetic insight into both partners. Here, we conduct the largest phylogenetic analysis of sea anemones (Order Actiniaria) to date, with a focus on expanding the biogeographic and taxonomic sampling of the 10 nominal clownfish-hosting species. Using a combination of mtDNA and nuDNA loci we test 1) the monophyly of each clownfish-hosting family and genus, 2) the current anemone taxonomy that suggests symbioses with clownfishes evolved multiple times within Actiniaria, and 3) whether, like the clownfishes, there is evidence that host anemones have a Coral Triangle biogeographic origin. Our phylogenetic reconstruction demonstrates widespread poly-and para-phyly at the family and genus level, particularly within the family Stichodactylidae and genus *Sticodactyla*, and suggests that symbioses with clownfishes evolved minimally three times within sea anemones. We further recover evidence for a Tethyan biogeographic origin for some clades. Our data provide the first evidence that clownfish and some sea anemone hosts have different biogeographic origins, and that there may be cryptic species of host anemones. Finally, our findings reflect the need for a major taxonomic revision of the clownfish-hosting sea anemones.

## 1. Introduction

Symbiosis often confers novel abilities or characteristics in at least one partner, can lead to adaptive radiation, and contributes meaningfully to the biodiversity within ecosystems. The clownfish-sea anemone symbiosis is an icon of tropical coral reefs of the Indo-West Pacific and is perhaps the most recognizable and famous example of symbiosis on the planet. The complexity of the clownfish-sea anemone symbiosis has attracted a great deal of popular and scientific attention and has been used as a model system to explore adaptive radiation, mutualism, specialism versus generalism, micro-and macro-evolution, animal behavior, social group structure and population dynamics, competition, venom resistance, host choice, larval dispersal and recruitment, biogeography, sex determination, and climate change among others (e.g. Almany et al., 2007; Beldade et al., 2017; Buston, 2004; Buston et al., 2007; Camp et al., 2016; Casas et al., 2018; Fautin, 1991; Hayashi et al., 2019; Huebner et al., 2012; Litsios & Salamin, 2014; Litsios et al., 2012, 2014a, 2014b; Mebs, 2009; Miyagawa-Kohshima et al., 2014; Ollerton et al, 2007; Schmitt & Holbrook, 2003; Szczebak et al., 2013). The charismatic nature of the relationship between anemones and clownfishes have made both constituents among the most heavily collected and sought after animals in the ornamental aquarium trade and species of conservation concern (Jones et al., 2008; Scott et al., 2014; Shuman et al., 2005; Rhyne et al., 2017; Wabnitz, 2003).

Our evolutionary understanding of this symbiosis, however, comes almost entirely from studies of clownfishes. Currently, there are 30 described species of clownfishes (or anemonefishes), which form the reciprocally monophyletic subfamily Amphiprioninae of the damselfish family Pomacentridae (e.g. Cooper et al., 2009; Litsios et al., 2012, 2014a; Rolland et al., 2018). Mutualism with sea anemones is believed to have been present in the common ancestor of all clownfishes (Litsios et al., 2012), which is estimated to have evolved ~12 mya in the Coral Triangle. The majority of clownfish diversity is the result of a recent adaptive radiation to a symbiotic lifestyle, with 25 of the 30 species having evolved within the last 5 mya (Litsios et al., 2012). Host specificities of clownfishes to sea anemones are also well resolved and span the host specialist-generalist continuum (e.g. Fautin, 1986; Fautin, 1991; Fautin & Allen, 1992; Litsios et al., 2012; Litsios et al., 2014a). Clownfish morphology and patterns of host specificity support the hypothesis that clownfishes have adapted to ecological niches associated with anemone hosts (Litsios et al., 2012).

Unlike the clownfishes, their sea anemone hosts have been poorly represented in systematic and phylogenetic studies, obscuring their biogeographic origin and patterns of diversification that have bearing on our interpretation of how the symbiosis has evolved. Broadly, sea anemones (Cnidaria: Anthozoa: Actiniaria) are a diverse group of benthic anthozoans that are found in every major marine habitat. Contrary to the generally observed pattern of hyperdiversity in the tropics, anemone diversity peaks in temperate marine ecosystems, where they often constitute the dominant benthic macrofauna (Fautin et al., 2013). The ecological success of sea anemones can be partly attributed to the diverse symbioses they form. This is particularly true for the clownfish-hosting anemones, which receive protection, nitrogen (via fish excrement), and increased gas transfer from their clownfish symbionts (Cleveland et al., 2011; Dunn, 1981; Roopin et al., 2008; Szczbak et al., 2013).

According to Fautin (2013), there are 10 species of sea anemone hosts distributed throughout the range of the clownfish symbiosis, a span that encompasses coral reef habitats from the Northern Red Sea through the Central Pacific Ocean (Dunn, 1981; Fautin, 1991; Fautin & Allen, 1992). Eight of the 10 nominal anemone species have widespread distributions and broadly overlapping biogeographic ranges (Fautin & Allen, 1992). All belong to the anemone superfamily Actinioidea, but are not taxonomically described to belong to a reciprocally monophyletic clade with a common ancestor that was symbiotic with clownfishes (Dunn, 1981; Fautin, 1991; Fautin & Allen, 1992). The 10 host species are presently interpreted to belong to three families (Actiniidae, Stichodactylidae, and Thalassianthidae) and encompass five genera (*Cryptodendrum*, *Entacmaea*, *Heteractis*, *Macrodactyla*, and *Stichodactyla*). The genera *Cryptodendrum* and *Entacmaea* are monospecific, while four host species are described as belonging to *Heteractis* and three to *Stichodactyla* (reviewed by Dunn, 1981). Two species are described to belong to the genus *Macrodactyla*, with only *M. doreensis* hosting clownfish.

Some disagreement exists regarding the status of the genus *Heteractis,* and there is confusion regarding the familial assignment of *Entacmaea*. England (1988) argued that because the type specimen for the genus *Heteractis* (*H. aurora*) harbors macrobasic amastigophore nematocysts (*p*-mastigophores A with looped tubule *sensu* Gusmão et al., 2018), this excludes *H. crispa, H. magnifica,* and *H. malu* from the genus. England (1998) thus resurrected *Radianthus* as a valid genus and reinstated the family Heteractidae to include *H. aurora*, *R. crispa, R. magnifica*, and *R. malu*, removing these two genera from the family Stichodactylidae. England’s work on the taxonomy of *Heteractis* was ignored for years and is not currently recognized in commonly used databases (e.g. Fautin 2013, WoRMS). Further, England (1987) considered and listed *E. quadricolor* as belonging to the family Stichodactylidae (rather than Actiniidae), noting that the basitrichs of the tentacles and column are different in Actiniidae and Stichodactylidae. England (1987) made no conclusion regarding the placement of *E. quadricolor* and thus it remains widely accepted as part of the family Actiinidae. Finally, an additional taxonomic issue with the clownfish-hosting anemones is that the generic sea anemone name *Macrodactyla* Haddon, 1898 is a junior homonym of a coleopteran genus (*Macrodactyla* Harris in Hitchcock, 1833), and the two anemone species included have been referred as belonging within the actiniid genus *Condylactis*, although without discussion (see Fautin, 2016).

To date, clownfish-hosting anemone species have only been described morphologically and have not been subjected to extensive molecular investigation. Anemones have simple body plans, no hard parts, and few diagnostic morphological characters, making traditional taxonomic descriptions challenging. Extensive phenotypic variation in living specimens makes field identification challenging, often making species identifications in the published literature unreliable (Dunn, 1981). Historically, this has led to widespread confusion regarding how many species of host anemones there actually are, and an abundance of species descriptions. For example, there are over 60 synonymized names for *E. quadricolor* alone in the World Registry of Marine Species (Daly & Fautin, 2019). These issues with traditional actiniarian taxonomy have rendered many non-clownfish hosting anemone genera para-or polyphyletic upon molecular phylogenetic investigation (e.g. Daly et al., 2017; Rodríguez et al., 2014). However, no phylogenetic study has included representatives from each of the 10 nominal host species (e.g. Daly et al., 2008, 2017; Rodríguez et al., 2014), and species that have been examined were often limited to a single individual sample per species, leaving their broader phylogenetic placement, taxonomy, and biogeography untested. Based on the currently accepted sea anemone taxonomy, it is expected that symbiosis with clownfishes evolved independently at least three times— once in each of the three families in which clownfish-host anemones are currently described. However, if England’s (1988) re-description of the family Heteractidae is supported, symbiosis with clownfishes would likely be expected to have evolved a fourth time, and possibly a fifth if *Entacmaea* does not belong within Actiinidae. Thus, thorough sampling and sequencing efforts are critical to shed light on the evolutionary history of these anemones and the clownfish symbiosis broadly, as well as to provide a more comprehensive understanding of actinarian diversity and evolution. Here, we conducted the largest phylogenetic analysis of Actiniaria to date, and tested 1) the monophyly of each clownfish-hosting family and genus, 2) the current anemone taxonomy that suggests symbiosis with clownfishes evolved multiple times within Actiniaria, and 3) examined if the clownfish hosting anemones, like their clownfish symbionts, have a biogeographic origin in the Coral Triangle.

## 2. Methods

### 2.1. Taxonomic sampling

Representatives from each of the 10 nominally described clownfish hosting sea anemones were included in this study (Figure 1). Data were acquired from a combination of field collected tissue samples, museum holdings, the aquarium trade, and GenBank (Table 1). Field collected tissue samples were collected by hand using SCUBA and preserved in 95% EtOH or RNAlater. We included a total of 59 individual samples across the 10 clownfish hosting anemone species, 43 of which were newly acquired for this study (Table 1). Where possible, we included samples from both the Indian and Pacific Ocean basins for nominal clownfish hosting anemones. Sampling effort was most exhaustive for *E. quadricolor* and *H. crispa* and we include multiple sample localities across their geographic ranges (Table 1). We also generated new sequence data for *S. helianthus*, a non-clownfish hosting sea anemone and the only member of the genus *Stichodactyla* native to the Tropical Western Atlantic (Table 1). Additional actinioidean in-group samples were obtained from previously published datasets (e.g. Daly et al., 2017; Rodríguez et al., 2014; Table S1). In total our analyses included 157 individuals from 91 species across the superfamily Actinioidea (Table1; Table S1). We further included 89 species representing the other four anemone superfamilies Metridioidea, Actinostoloidea, Edwardsioidea, and Actinernoidea (Table S1). Two non-actiniarian samples from Corallimorpharia and Zoantharia were included as outgroups (Table S1). The final dataset of 256 individuals across 180 anemone species represents the largest phylogenetic analysis of actiniarian diversity to date.

**Table 1.** Clownfish-hosting sea anemone species and samples included in this study as part of the broader phylogenetic analysis of Actiniaria. Table includes information on Family, Genus, and Species designations based on the currently accepted taxonomy from Fautin (2013). All species listed belong to superfamily Actinioidea. Locality indicates country of origin for each sample, and Ocean Basin reflects the body of water each locality broadly belongs to. GenBank accession numbers in **Bold** are new to this study. *Denotes *Stichodactyla helianthus* from the Tropical Western Atlantic-a non-clownfish host species.

| Family | Genus | Species | Locality | Ocean Basin | CO3 | 12S | 16S | 18S | 28S |
| --- | --- | --- | --- | --- | --- | --- | --- | --- | --- |
| Actiniidae | *Entacmaea* | *quadricolor* | Japan | West Pacific | **MK522439** | **MK519401** | **MK519456** | - | - |
| Actiniidae | *Entacmaea* | *quadricolor* | Japan | West Pacific | **MK522440** | **MK519402** | **MK519457** | **MK519571** | - |
| Actiniidae | *Entacmaea* | *quadricolor* | Maldives | Central Indian | **MK522441** | **MK519403** | - | **MK519566** | **MK519639** |
| Actiniidae | *Entacmaea* | *quadricolor* | Maldives | Central Indian | **MK522442** | **MK519404** | **MK519458** | **MK519567** | **MK519644** |
| Actiniidae | *Entacmaea* | *quadricolor* | Saudi Arabia | Red Sea | **MK522443** | **MK519405** | **MK519459** | **MK519568** | **MK519643** |
| Actiniidae | *Entacmaea* | *quadricolor* | Tonga | Southwest Pacific | **MK522444** | **MK519406** | **MK519460** | **MK519569** | **MK519645** |
| Actiniidae | *Entacmaea* | *quadricolor* | Tonga | Southwest Pacific | **MK522445** | **MK519407** | **MK519461** | **MK519570** | **MK519646** |
| Actiniidae | *Entacmaea* | *quadricolor* | United Arab Emirates | Gulf of Oman | **MK522446** | **MK519408** | **MK519462** | **MK519572** | **MK519640** |
| Actiniidae | *Entacmaea* | *quadricolor* | United Arab Emirates | Gulf of Oman | **MK522447** | **MK519409** | **MK519463** | **MK519573** | **MK519641** |
| Actiniidae | *Entacmaea* | *quadricolor* | United Arab Emirates | Gulf of Oman | **MK522448** | **MK519410** | **MK519464** | - | **MK519642** |
| Actiniidae | *Macrodactyla* | *dorensis* | Philippines | Coral Triangle | **MK522449** | **MK519411** | **MK519465** | **MK519581** | **MK519661** |
| Actiniidae | *Macrodactyla* | *dorensis* | Philippines | Coral Triangle | **MK522450** | - | **MK519466** | **MK519582** | - |
| Actiniidae | *Macrodactyla* | *dorensis* | Unknown | Unknown | GU473342 | EU190739 | EU190785 |  | KJ483049 |
| Stichodactylidae | *Heteractis* | *aurora* | Maldives | Central Indian | **MK522451** | **MK519412** | **MK519467** | - | - |
| Stichodactylidae | *Heteractis* | *aurora* | Maldives | Central Indian | **MK522452** | **MK519413** | **MK519468** | - | - |
| Stichodactylidae | *Heteractis* | *aurora* | Maldives | Central Indian | **MK522453** | **MK519414** | **MK519469** | - | **MK519647** |
| Stichodactylidae | *Heteractis* | *aurora* | Maldives | Central Indian | **MK522454** | **MK519415** | **MK519470** | - | - |
| Stichodactylidae | *Heteractis* | *aurora* | Unknown | Unknown | KC812229 | KC812139 | KC812160 | KC812184 | - |
| Stichodactylidae | *Heteractis* | *aurora* | Unknown | Unknown | - | EU190729 | EU190773 | - | - |
| Stichodactylidae | *Heteractis* | *aurora* | Unknown | Unknown | - | - | - | KC812185 | - |
| Stichodactylidae | *Heteractis* | *crispa* | Unknown | Unknown | KC812230 | KC812140 | KC812161 | KC812186 | - |
| Stichodactylidae | *Heteractis* | *crispa* | Japan | West Pacific | **MK522455** | **MK519416** | **MK519471** | - | **MK519648** |
| Stichodactylidae | *Heteractis* | *crispa* | Japan | West Pacific | **MK522456** | **MK519417** | **MK519472** | - | **MK519649** |
| Stichodactylidae | *Heteractis* | *crispa* | Japan | West Pacific | **MK522457** | **MK519418** | **MK519473** | - | **MK519650** |
| Stichodactylidae | *Heteractis* | *crispa* | Palau | West Pacific | **MK522458** | **MK519419** | **MK519474** | - | - |
| Stichodactylidae | *Heteractis* | *crispa* | Palau | West Pacific | **MK522460** | **MK519421** | **MK519476** | - | - |
| Stichodactylidae | *Heteractis* | *crispa* | Palau | West Pacific | **MK522461** | **MK519422** | **MK519477** | - | - |
| Stichodactylidae | *Heteractis* | *crispa* | Saudi Arabia | Red Sea | **MK522462** | **MK519423** | **MK519478** | - | **MK519654** |
| Stichodactylidae | *Heteractis* | *crispa* | Saudi Arabia | Red Sea | **MK522463** | **MK519424** | **MK519479** | - | - |
| Stichodactylidae | *Heteractis* | *crispa* | Saudi Arabia | Red Sea | **MK522464** | **MK519425** | **MK519480** | - | - |
| Stichodactylidae | *Heteractis* | *crispa* | Tonga | Southwest Pacific | **MK522465** | **MK519426** | **MK519481** | **MK519574** | **MK519651** |
| Stichodactylidae | *Heteractis* | *crispa* | United Arab Emirates | Gulf of Oman | **MK522466** | **MK519427** | **MK519482** | - | - |
| Stichodactylidae | *Heteractis* | *crispa* | United Arab Emirates | Gulf of Oman | **MK522467** | **MK519428** | **MK519483** | - | **MK519652** |
| Stichodactylidae | *Heteractis* | *crispa* | United Arab Emirates | Gulf of Oman | **MK522468** | **MK519429** | **MK519484** | - | **MK519653** |
| Stichodactylidae | *Heteractis* | *magnifica* | Unknown | Unknown | KJ482988 | EU190732 | EU190777 | - | JK483093 |
| Stichodactylidae | *Heteractis* | *magnifica* | Unknown | Unknown | GU473361 | - | - | EU190862 | EU190821 |
| Stichodactylidae | *Heteractis* | *magnifica* | Unknown | Unknown | KC812231 | - | KC812162 | KC812187 | - |
| Stichodactylidae | *Heteractis* | *magnifica* | Maldives | Central Indian | **MK522469** | **MK519430** | **MK519485** | **MK519575** | **MK519656** |
| Stichodactylidae | *Heteractis* | *magnifica* | Maldives | Central Indian | **MK522470** | **MK519431** | **MK519486** | **MK519576** | **MK519657** |
| Stichodactylidae | *Heteractis* | *magnifica* | Maldives | Central Indian | **MK522471** | **MK519432** | **MK519487** | **MK519577** | **MK519658** |
| Stichodactylidae | *Heteractis* | *magnifica* | Maldives | Central Indian | **MK522472** | **MK519433** | **MK519488** | **MK519578** | **MK519659** |
| Stichodactylidae | *Heteractis* | *magnifica* | Maldives | Central Indian | **MK522473** | **MK519434** | **MK519489** | **MK519579** | **MK519660** |
| Stichodactylidae | *Heteractis* | *magnifica* | Palau | West Pacific | **MK522459** | **MK519420** | **MK519475** | **MK519580** | **MK519655** |
| Stichodactylidae | *Heteractis* | *malu* | Tonga | Southwest Pacific | **MK522474** | - | - | - | - |
| Stichodactylidae | *Heteractis* | *malu* | Tonga | Southwest Pacific | **MK522475** | - | **MK519490** | - | - |
| Stichodactylidae | *Stichodactyla* | *gigantea* | Unknown | Unknown | GU473347 | EU190747 | - | EU190873 | EU190835 |
| Stichodactylidae | *Stichodactyla* | *gigantea* | Unknown | Unknown | KC812232 | - | - | KC812188 | - |
| Stichodactylidae | *Stichodactyla* | *gigantea* | Unknown | Unknown | KY789299 | - | EU190793 | - | - |
| Stichodactylidae | *Stichodactyla* | *haddoni* | Philippines | Coral Triangle | **MK522476** | **MK519435** | **MK519491** | **MK519583** | **MK519662** |
| Stichodactylidae | *Stichodactyla* | *haddoni* | Unknown | Unknown | KC812233 | - | FJ417090 | FJ417089 | - |
| Stichodactylidae | *Stichodactyla* | *helianthus\** | Panama | Tropical Western Atlantic | **MK522477** | **MK519436** | **MK519492** | **MK519563** | **MK519667** |
| Stichodactylidae | *Stichodactyla* | *helianthus\** | Panama | Tropical Western Atlantic | **MK522478** | **MK519437** | **MK519493** | **MK519564** | **MK519668** |
| Stichodactylidae | *Stichodactyla* | *helianthus\** | Panama | Tropical Western Atlantic | **MK522479** | **MK519438** | **MK519494** | - | **MK519669** |
| Stichodactylidae | *Stichodactyla* | *helianthus\** | Panama | Tropical Western Atlantic | **MK522480** | **MK519439** | **MK519495** | - | **MK519670** |
| Stichodactylidae | *Stichodactyla* | *mertensii* | Unknown | Tropical Western Atlantic | KC812234 | KC812141 | KC812163 | KC812189 | - |
| Stichodactylidae | *Stichodactyla* | *mertensii* | Maldives | Central Indian | **K522481** | **MK519440** | **MK519496** | **MK519584** | **MK519663** |
| Stichodactylidae | *Stichodactyla* | *mertensii* | Maldives | Central Indian | **K522482** | **MK519441** | **MK519497** | **MK519585** | **MK519664** |
| Stichodactylidae | *Stichodactyla* | *mertensii* | Maldives | Central Indian | **K522483** | **MK519442** | **MK519498** | **MK519586** | **MK519665** |
| Stichodactylidae | *Stichodactyla* | *mertensii* | Maldives | Central Indian | - | **MK519443** | **MK519499** | **MK519587** | **MK519666** |
| Thalassianthidae | *Cryptodendrum* | *adhaesivum* | Maldives | Central Indian | **K522484** | -- | **MK519500** | **MK519565** | **MK519638** |
| Thalassianthidae | *Cryptodendrum* | *adhaesivum* | Unknown | Unknown | KC812235 | KC812142 | KC812163 | KC812190 | KC812214 |
| Thalassianthidae | *Cryptodendrum* | *adhaesivum* | Unknown | Unknown | KC812236 | KC812143 | KC812165 | KC812191 | KC812215 |
| Thalassianthidae | *Cryptodendrum* | *adhaesivum* | Unknown | Unknown | KC812237 | KC812144 | KC812166 | KC812192 | KC812216 |

**Figure 1.**
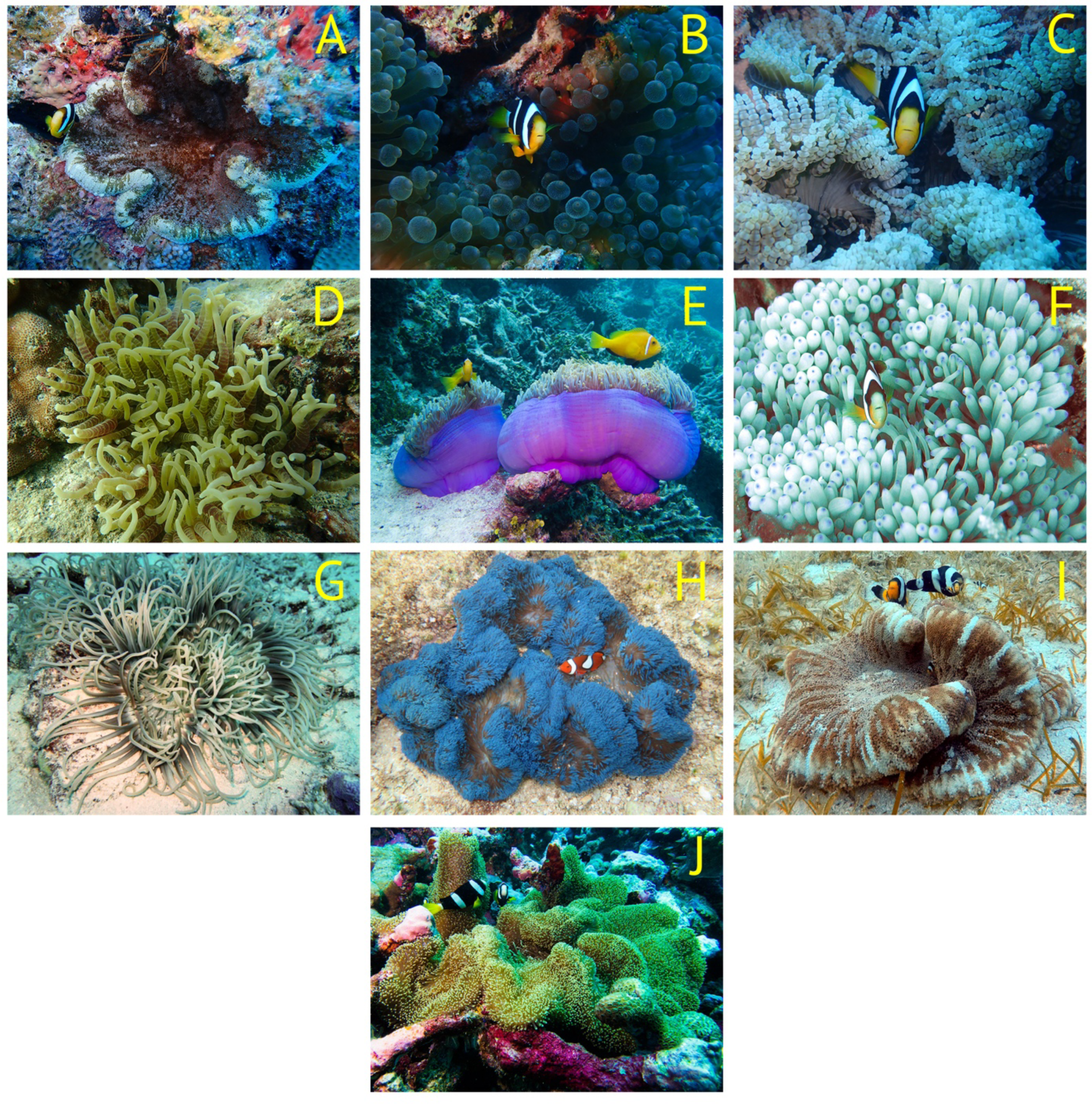
Representative images of the 10 species of clownfish-hosting sea anemones. A) *Cryptodendrum adhaesivum*, B) *Entacmaea quadricolor*, C) *Heteractis aurora*, D) *Heteractis crispa*, E) *Heteractis magnifica*, F) *Heteractis malu* (photo credit: Matthew Lee), G) *Macrodactyla doreensis*, H) *Stichodactyla gigantea* (photo credit: Anne Hoggett), I) *Stichodactyla haddoni* (photo credit: Singgih Afifa Putra), J) *Stichodactyla mertensii*. Photo of *M. doreensis* by Lyle Vail, used under creative commons license attribution 3.0 unported license http://creativecommons.org/licenses/by/3.0/. Photos A-E, and J by B. Titus.

### 2.2. DNA extraction, PCR, and sequencing

Genomic DNA was extracted using the DNeasy Blood and Tissue Kits (QIAGEN Inc.) and stored at −20ºC. All DNA extractions were standardized to ~20ng/µL, and template DNA was amplified from genomic samples using published primers and standard PCR techniques. We targeted three mitochondrial (partial 12S rDNA, 16S rDNA, and CO3) and two nuclear (18S rDNA, and partial 28S rDNA) gene markers for phylogenetic reconstruction. Primer sequences and PCR run conditions can be found elsewhere (Daly et al., 2008; Geller & Walton 2001; Gusmão & Daly, 2010). All PCR products were cleaned using ExoSAP-IT (Thermo Fisher) and FastAP (Thermo Fisher) enzyme reactions. Each PCR clean up used 10 µL PCR product, 2 µL ExoSAP-IT, and 1 µL FastAP. Clean up reactions were carried out under the following conditions: 37ºC for 15 min, 85ºC for 15 min, and hold at 4ºC. Cycle sequencing reactions were carried out using 5 µL purified PCR product at a concentration of 25 ng of PCR product for every 200 base pairs of length. Cycle sequence products were cleaned using the Sephadex G-50 (Sigma-Aldrich) spin-column protocol. Samples were sequenced using traditional Sanger-based capillary electrophoresis on an ABI 3730 at the American Museum of Natural History (AMNH). Forward and reverse sequences were assembled in Sequencher v. 4.9 (Gene Codes Corporation, Ann Arbor, MI) and compared (via BLAST) against the nucleotide database of GenBank to determine whether the target organism was sequenced, rather than an endosymbiotic dinoflagellate. Newly generated sequences have been deposited in GenBank (Table 1; Table S1).

### 2.3. Phylogenetic analyses

Newly assembled sequences were aligned for each locus separately using MAFFT v7.394 (Katoh & Standley, 2013) on the CIPRES Science Gateway Portal (Miller et al., 2010) under the following alignment parameters: Strategy, L-INS-i; Scoring matrix for nucleotide sequences, 200PAM/k=2; Gap opening penalty, 1.53; Offset value, 0.1; Max iterate, 1000; Retree, 1. Additionally, multiple sequence alignments for each locus were analyzed using Gblocks v0.91 (Castresana, 2000) to remove poorly aligned and/or mutationally saturated divergent regions. The following parameters were used in the Gblocks: Maximum number of contiguous non-conserved positions, 8; Minimum length of a block, 5; Gap positions allowed. Gblocks analysis for the 28s gene further included: Allow smaller final blocks; Allow less strict flanking positions. The complete Gblocks sequence alignments for each locus were concatenated into a final super matrix and analyzed using PartitionFinder2 (Lanfear et al., 2016) to identify the best partitioning scheme and best fit model of nucleotide evolution under the corrected Akaike Information Criterion (AIC).

Maximum Likelihood (ML) phylogenetic analyses were conducted using RAxML v8.2.10 as implemented on the CIPRES Science Gateway Portal. Analyses were conducted using four partitions as determined by PartitionFinder: CO3, 12S/16S, 18S, and 28S. For each partition we implemented a GTR+G+I model of nucleotide substitution as determined by PartitionFinder analyses. Clade support was analysed using rapid bootstrapping with a subsequent ML search, and we let RAxML halt bootstrapping automatically using MRE-based bootstrapping criterion. RAxML analyses recovered the 10 nominal clownfish hosting species to belong to three clades (see Results). However, poor nodal support across the backbone of the superfamily Actinioidea could not resolve the relationship among these clades and other clades within Actinioidea, and thus, was unable to distinguish between one and three independent evolutionary origins of symbiosis with clownfishes. Thus, we conducted a likelihood ratio test of tree topologies (SH test) between the initial unconstrained analysis above, and a separate analysis where the 10 clownfish hosting species were constrained to a monophyletic group within a monophyletic Actinioidea. The SH test was conducted using RAxML on the CIPRES Science Gateway Portal.

Finally, we conducted separate RAxML analyses for each anemone clade in which our larger Actiniaria-wide analyses inferred an independent evolutionary origin of symbiosis with clownfishes (see Results). These analyses were conducted at shallower evolutionary levels in order to include all sequence data that may have otherwise been excluded by Gblocks in our larger dataset in an attempt to improve phylogenetic resolution within each group and explore potential biogeographic signal. Data were concatenated, partitioned, and analyzed in RAxML as above. The three analyses included 1) all members of the family Thalassianthidae, genus *Stichodactyla*, and *H. magnifica*, 2) all members of *H. aurora, H. crispa, H. malu*, and *M. doorenensis*, and 3) all members of *E. quadricolor*.

## 3. Results

Concatenated Gblocks alignments resulted in a final data matrix of 5,887 base pairs for the full actiniarian-wide analysis. Alignments for lower level analyses of 1) all members of the family Thalassianthidae, genus *Stichodactyla*, and *H. magnifica*, 2) all members of *H. aurora, H. crispa, H. malu*, and *M. doorenensis,* and 3) *E. quadricolor* resulted in data matrices of 6,941, 7,085, and 6,871 base pairs respectively.

Maximum Likelihood phylogenetic analyses in RAxML recovered each anemone superfamily, and hierarchical relationships among superfamilies, as described by Rodríguez et al (2014), but with varying degrees of nodal support (Figure S1). Like Rodríguez et al. (2014), we obtained inconsistent nodal support across the backbone of Actiniaria, which is common with this suite of phylogenetic markers. Relationships within superfamilies Actinernoidea, Actinostoloidea, Edwardsioidea, and Metridioidea were also broadly reflective of those recovered by Rodríguez et al. (2014; Figure S2).

Within Actinioidea, our ML analyses recovered the family Stichodactylidae (as currently accepted, including *Stichodactyla* and *Heteractis*), as polyphyletic (Figure 2; Figure S3). Only the magnificent anemone *H. magnifica* was recovered in the clade that included members of the genus *Stichodactyla*. Our analyses also find the genus *Stichodactyla* to be paraphyletic (Figure 2; Figure S3). In our full analysis, specimens of *S. mertensii* and *S. haddoni* appeared to share a more recent common ancestor with *C. adhaesivum*, *Thalassianthus aster,* and *T. hemprichii* than with *S. gigantea* and *S. helianthus* (Figure 2; Figure S3). Our in-depth ML analysis of this clade did not recover this relationship but instead reflected that all species from the Indo-Pacific (*S. gigantea*, *S. haddoni*, *S. mertensi*, *H. magnifica*, *C. adhaesivum*, and the genus *Thalassianthus*) are more closely related to each other than they are to the Atlantic *S. helianthus* (Figure 3). Interestingly, our analyses recovered the family Thalassianthidae, which includes *Cryptodendrum* and *Thalassianthus*, to be nested within a well-supported clade that included all members of the genus *Stichodactyla* and *H. magnifica* (Figures. 2 & 3; Figure S3). We refer to this clade as “Stichodactylina” or “true carpet anemones” throughout the remainder of the manuscript. The species *S. helianthus* appears to have diverged early in the evolutionary history of Stichodactylina, which split the clade into Atlantic and Indo-Pacific sister lineages (Figures 2 & 3; Figure S3).

**Figure 2.**
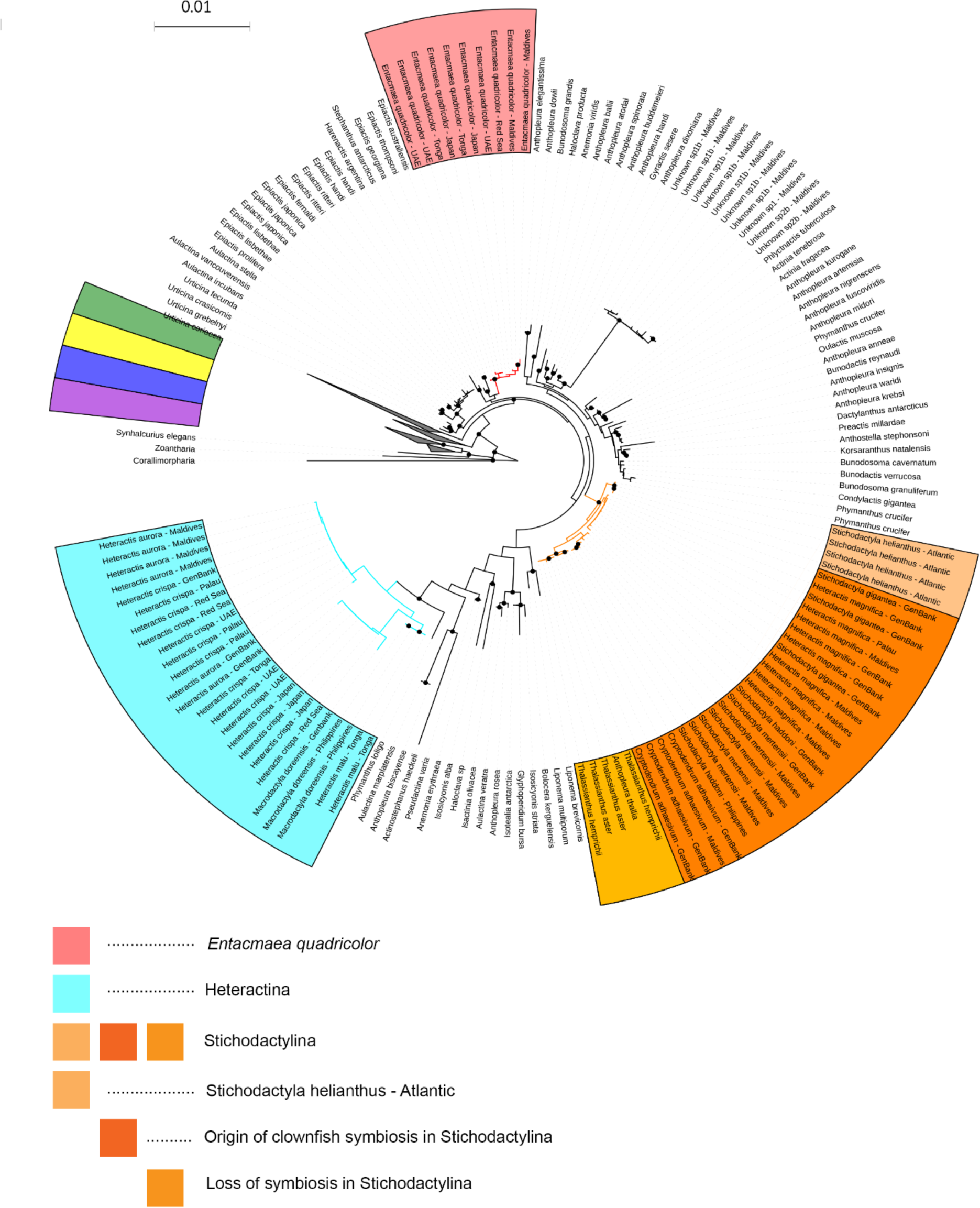
Phylogenetic reconstruction of Actiniaria. Tree resulting from Maximum Likelihood (ML) analysis in RAxML of concatenated dataset (CO3, 12S, 16S, 18S, 28S). Tree reflects relationships within anemone superfamily Actinioidea, with remaining superfamilies Actinernoidea (purple), Actinostoloidea (yellow), Edwardsioidea (blue), and Metridioidea (green) collapsed for clarity. Legend highlights the three clades where symbiosis with clownfishes has evolved: Red = *Entacmaea quadricolor*, Light Blue = Heteractina (*Heteractis aurora*, *H. crispa*, *H. malu*, and *Macrodactyla doreensis*), Oranges = Stichodactylina (*Cryptodendrum adhaesivum, H. magnifica, Stichodactyla gigantea, S. haddoni, S. helianthus, S. mertensii, Thalassianthus aster, T. hemprichii*). Within Stichodactylina different hues of orange represent where *S. helianthus* diverged into the Atlantic Ocean, where symbiosis with clownfishes arose in the Indo-West Pacific, and where symbiosis with clownfishes may have been lost in the genus *Thalassianthus*. Black filled circles represent nodes with bootstrap resampling values ≥ 70.

**Figure 3.**
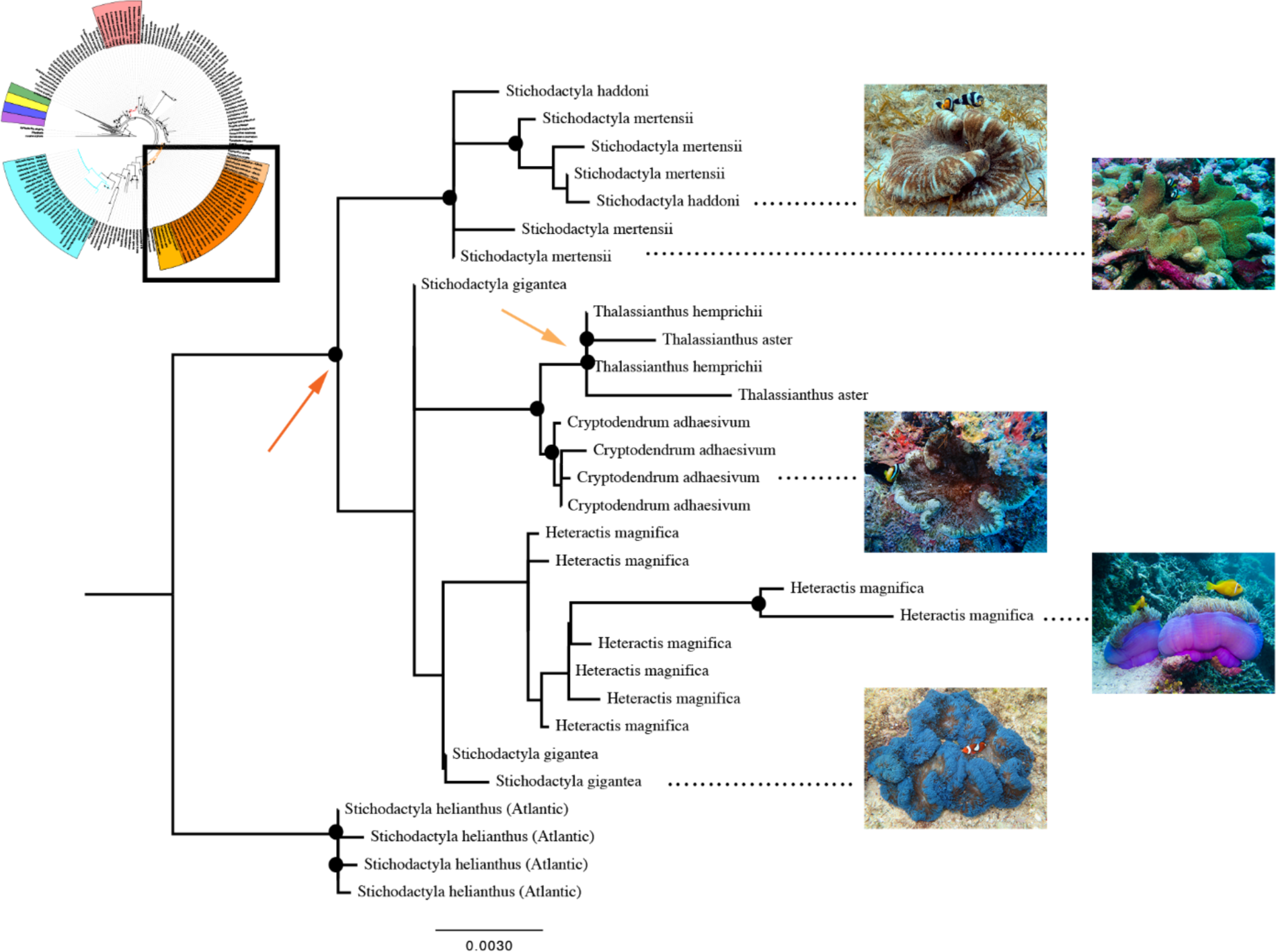
Partitioned Maximum Likelihood phylogenetic reconstruction of the carpet sea anemone Stichodactylina clade using all available sequence data (CO3; 12S, 16S, 18S, 28S). Orange arrow represents potential origin of symbiosis with clownfishes. Light orange arrow represents potential loss of symbiosis with clownfishes in the genus *Thalassianthus*. Colored nodes reflect bootstrap resampling values ≥75. Pictured are the five host species *C<u>r</u>yptodendrum adhaesivum, Heteractis magnifica, Stichodactyla gigantea, S. haddoni,* and *S. mertensii*.

Other than *H. magnifica*, the species currently ascribed to the genus *Heteractis sensu* Dunn (1981) (or to *Heteractis* and *Radianthus sensu* England (1988)), formed a moderately supported clade, along with *M. doreensis*. This clade is within the superfamily Actinioidea and distant to the Stichodactylina (Figure 2; Figure S3). This group of *Heteractis* and *Macrodactyla* species shared a well-supported node with *Phymanthus loligo* deeper in the tree (Figure 2 & 4; Figure S3). Unexpectedly, our analyses find three distinct groupings of individuals identified as *H. aurora, H. crispa, H. malu,* and *M. doreensis* separated by relatively long branches, but with little nodal support for those groupings. Further, these groupings do not correspond to any current sea anemone taxonomy, as individuals identified as members of each nominal species were found within each cluster (Figure 2 & 4; Figure S3). Although this clade included *M. doreensis* and excluded *H. magnifica*, it does appear to partially support England’s (1988) distinction between these species and members of Stichodactylidae. We refer to this clade as “Heteractina”, in reference to England’s designation, throughout the rest of the manuscript. In-depth phylogenetic analyses of Heteractina did not provide further resolution among species in this clade (Figure 4).

**Figure 4.**
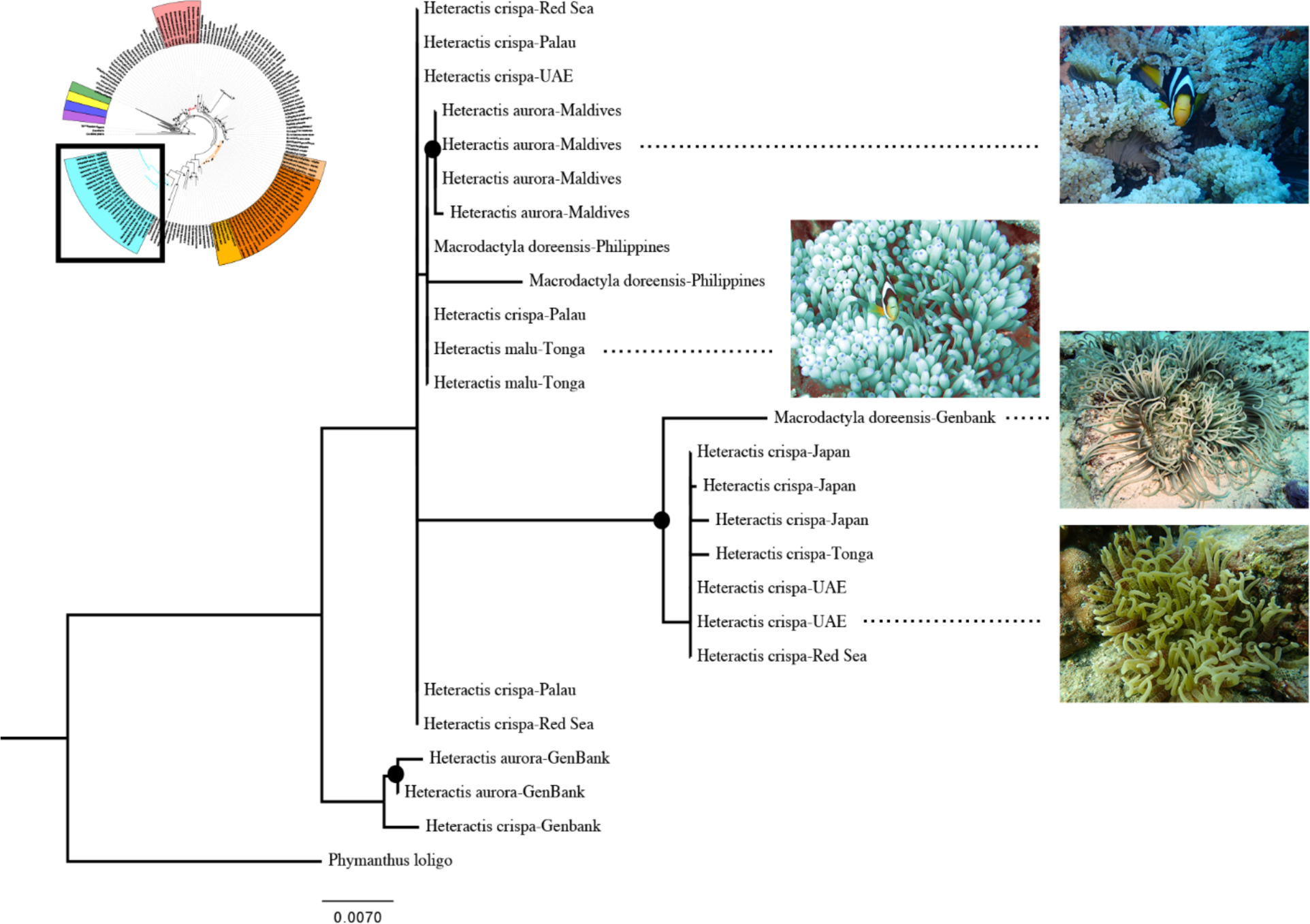
Partitioned Maximum Likelihood phylogenetic reconstruction of the Heteractina clade using all available sequence data (CO3; 12S, 16S, 18S, 28S). Colored nodes reflect bootstrap resampling values ≥75. Pictured are the four host anemone species *Heteractis aurora, H. crispa, H. malu,* and *Macrodactyla doreensis*.

The individuals of bubble tip anemone *E. quadricolor* formed a highly supported clade, which makes it, together with the adhesive anemone *C. adhaesivum*, one of only two clownfish hosting species to form well-supported monophyletic lineages in our actiniarian-wide analysis (Figure 2; Figure S3). No members of Actinioidea formed a well-supported sister relationship to *E. quadricolor,* although the relationship between *E. quadricolor* and a clade of Southern Hemisphere brooding anemones from the genus *Epiactis* would be worth further exploring with genomic data (Figure 2; Figure S3). Intraspecific branch lengths within *E. quadricolor* exceeded those typically found at the intraspecific levels in other species (Figure 2; Figure S3), and our shallow evolutionary analyses recovered well-supported hierarchical relationships that correspond to biogeographic patterns (Figure 5). We recovered two well-supported clades within *E. quadricolor*, one that encompasses samples from the Red Sea, Maldives, and Japan, and a second that includes samples from Tonga and Japan (Figure 5). Samples from the United Arab Emirates appear to have diverged early and likely represents a third lineage, although this node was not well supported (Figure 5).

**Figure 5.**
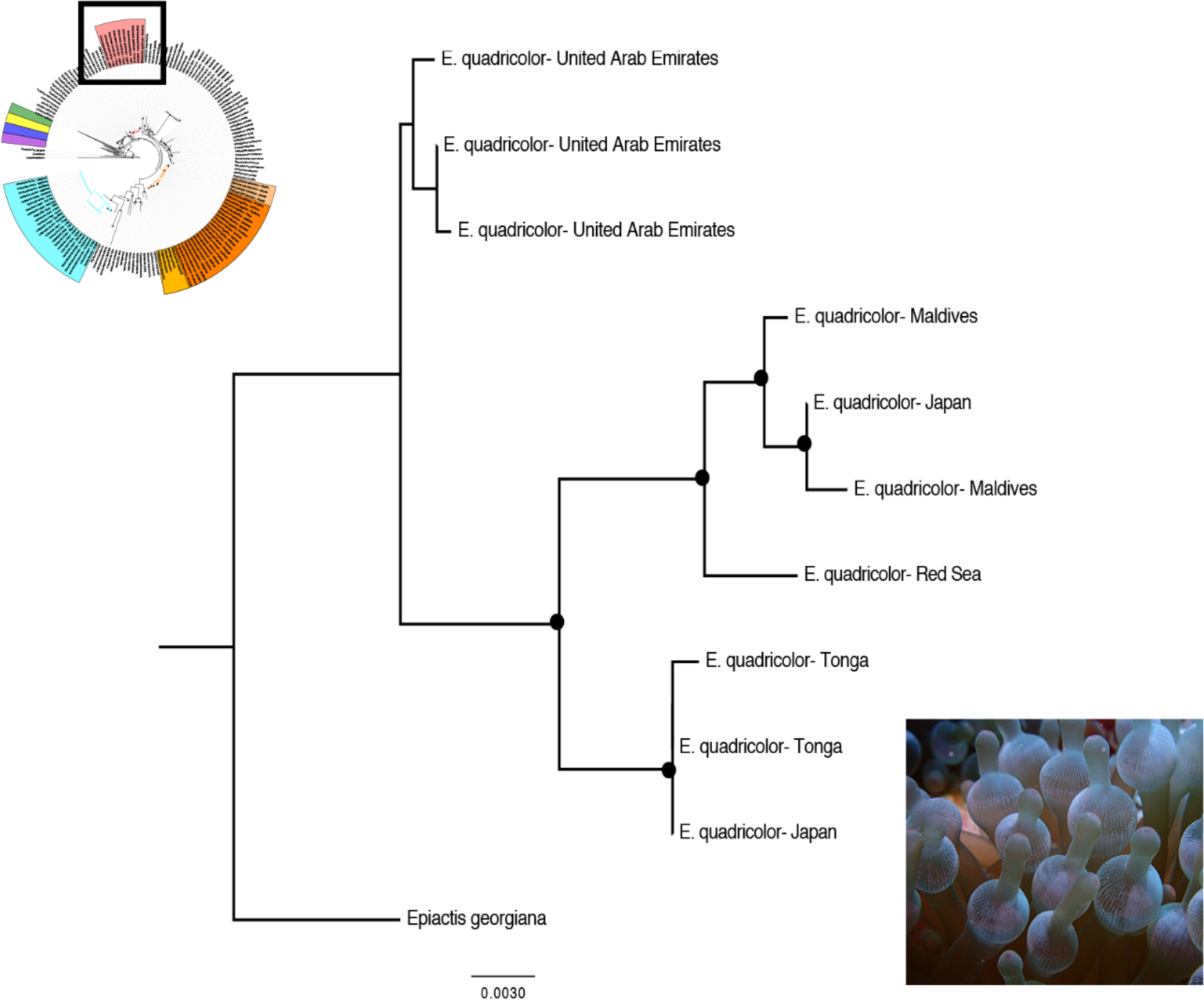
Partitioned Maximum Likelihood phylogenetic reconstruction of the bubble-tip sea anemone *Entacmaea quadricolor* using all available sequence data (CO3; 12S, 16S, 18S, 28S). Tree reflects samples collected from throughout the biogeographic range of the species. Colored nodes reflect bootstrap resampling values ≥75.

Likelihood ratio tests in RAxML show that our unconstrained analysis of Actiniaria is a significantly better topology than a topology that constrains all clownfish-hosting lineages to a monophyletic group [D(LH) = −709.74, p < 0.01]. Thus, based on our unconstrained Actiniaria-wide analysis, we conclude that symbiosis with clownfishes has evolved at least three times within sea anemones (Figure 2; Figure S3); once within the lineage currently construed as *E. quadricolor*, once within the true carpet anemones (Stichodactylina) after *S. helianthus* diverged in the Atlantic, and once within Heteractina (Figure 2; Figure S3). Based on our phylogenetic reconstruction, we identify the genus *Thalassianthus*, within Stichodactylina, as the only group where symbiosis with clownfishes has been lost (Figures 2 & 3; Figure S3).

## 4. Discussion

Our phylogenetic reconstruction of Actiniaria provides the first in-depth molecular investigation into the evolution of the clownfish-hosting sea anemones, shedding light on their systematics, diversity and biogeographic origin. This perspective has been lacking in broader studies of clownfish evolution (e.g. Litsios et al., 2014a). Perhaps unsurprisingly, we recover widespread poly-and para-phyly among the actiniarian families and genera to which the 10 host anemone species are assigned. Instead, we identify three higher-level clades (Stichodactylina, Heteractina, and *E. quadricolor*) where we infer symbioses with clownfishes have evolved independently. Below, we discuss the taxonomic problems our data have revealed at the family, generic, and species levels within the clownfish-hosting sea anemones. We follow with discussion of how our results impact our understanding of how clownfish-sea anemone symbiosis has evolved.

Family-level taxonomy within Actinioidea is messy, with no evidence of monophyly for most groups (Rodríguez et al. 2014; Larson & Daly 2016; Daly et al. 2017). The clownfish-hosting anemones are no exception to this general pattern. As outlined above, the clownfish hosting sea anemones are traditionally recognized by Fautin (1991, 2013, 2016) as belonging to three families. These include Actiniidae (*Entacmaea* and *Macrodactyla*), Stichodactylidae (*Stichodactyla* and *Heteractis*), and Thallasiandthidae (*Cryptodendrum*). England (1988) proposed four families, splitting *Heteractis* into Heteractidae, and further suggesting that Heteractidae includes *Heteractis,* with the single species *H. aurora,* and *Radianthus*, for the species commonly referred to as *H. crispa, H. magnifica,* and *H. malu*. Our molecular data provide some evidence for a clade that aligns with Stichodactylidae (our Stichodactylina) and for some aspects of the proposed Heteractidae, but the membership of these groups in our trees is not identical to their traditionally recognized taxonomic composition. Further, Stichodactylina also includes what is traditionally recognized as family Thalassianthidae (see Daly & Fautin 2019). In the broader phylogenetic picture, Actiniidae is paraphyletic, consisting of all Actinioidea not assigned to a more exclusive group.

The generic and familial taxonomic problems are most significant in *Heteractis* and *Stichodactyla*. Our molecular results do not support maintaining the current taxonomy of Fautin (1991) but are also inconsistent with the proposal of England (1988). In partial agreement with England (1988), we find that *H. aurora, H. crispa,* and *H. malu* do not from a monophyletic familial relationship with the members of the genus *Stichodactyla*, and thus partially support his proposal for the designation of a family Heteractidae. However, we find that *H. magnifica* is more closely related to species in *Stichodactyla* (and to *Cryptodendron* and *Thalassianthus*) than it is to the other species of *Heteractis.* Further, contra the proposal of England (1988) that it is distinct from the other species of the genus, *H. aurora* cannot be differentiated from *H. crispa* and *H. malu*, and thus, our molecular data do not support the resurrection of the genus *Radianthus* based on the presence of *p*-mastigophores A with a looped tubule in the cnidom of *H. aurora*. The nematocysts that were observed by England in *H. aurora* may be interpreted as being an immature stage of the *p*-mastigophores A present in all the species of *Heteractis* (Gusmão et al., 2018). Specimens of *M. doreensis* were also recovered to form a close relationship with *H. aurora*, *H. crispa*, and *H. malu*. This relationship between *Macrodactyla* and species of *Heteractis* contradicts the decision of Fautin (2016) to move these species to *Condylactis* to correct the nomenclature issue with the generic name *Macrodactyla*. *Macrodactyla* is quite distinct, phylogenetically, from the representative of *Condylactis* in our analyses (*C. gigantea*), which groups as sister to one Caribbean species of *Phymanthus* (Figure 2; Figure S3). Taken together, the relationships between *H. aurora*, *H. crispa*, *H. malu,* and *M. doreensis* form our higher-level Heteractina clade.

Our data also highlight generic and familial taxonomic problems of Fautin’s and England’s genus *Stichodactyla* and family Thalassianthidae. The nesting of the Thalassianthidae within *Stichodactyla*, coupled with the relationships among members of the genus *Stichodactyla* and *H. magnifica* renders *Stichodactyla* paraphyletic. *Heteractis magnifica* and all other Indo-Pacific members of *Stichodactyla, Cryptodendrum*, and *Thalassianthus* share a more recent common ancestor than they do with the Atlantic *S. helianthus*. *Stichodactyla mertensii* Brandt, 1835 is the type species of the genus *Stichodactyla* and thus, *S. helianthus* (Ellis, 1768) should likely be designated as belonging to a different genus according to the Principle of Priority (Art. 23, ICZN 1999). Similarly, *H. magnifica* should be placed in a different genus in concert with a revision of *Stichodactyla*. Together, all members of *Stichodactyla*, *Cryptodendrum, Thalassianthus*, and *H. magnifica* form our Stichodactylina clade.

At the species level, our data reveal that only *C. adhaesivum* and *E. quadricolor* were recovered as monophyletic with high support. Within our Heteractina clade, the pattern of relationships we find for the samples of *H. aurora*, *H. crispa, H. malu*, and *M. doreensis* are difficult to interpret. Although there appears to be a moderate amount of genetic variation within Heteractinia, the major nodes within the clade are all poorly supported and do not form species-specific units (Figure 4). It is possible that the repeated occurrence of subclades identified as *H. aurora* and *H. crispa* and *H. malu* suggest that these names encompass several as-yet undescribed species (Figure 4). Specimens of *M. doreensis* were also recovered in several of the clades within Heteractina. Two possible hypotheses may explain the patterns we see here: 1) *H. crispa, H. malu,* and *M. doreensis* are superficially similar in appearance (Figure 1), share similar habitat space, and could easily be misidentified in the field when these species co-occur. Multiple sequences in our analyses come from GenBank (Table 1) and thus it is possible that misidentification could have led to the patterns we observed here. We believe the likelihood for misidentification within our dataset is low, as the newly generated data for this study came from individuals that were collected, photographed in the field, and positively identified by the authors of this study directly. Although we identified anemones in the field following Fautin and Allen (1992), recent contributions to sea anemone systematics keep pointing to high levels of morphological convergence and the necessity of reevaluating the traditionally used morphological characters and their phylogenetic signal (e.g. Daly et al., 2017 Lauretta et al., 2014, Rodríguez et al., 2012, 2014). A more likely possibility is that 2) these genetic markers simply cannot resolve the relationship between these taxa at this level, and/or incomplete lineage sorting is driving the observed pattern among species. New, high resolution genomic methods, such as bait-capture approaches targeting ultra conserved elements (e.g. Quattrini et al., 2018), are needed to clarify these shallow relationships.

Within Stichodactylina we are unable to resolve species-level support for *S. gigantea, S. haddoni*, and *S. mertensii* (Figure 3). Samples of *S. haddoni* and *S. mertensii* do intermingle within a fairly well supported group but we do not recover species-level resolution. *Stichodactyla haddoni* and *S. mertensii* are described as occupying distinct ecological habitats, with *S. haddoni* occurring exclusively on sandy bottoms and *S. mertensii* occurring exclusively on coral-dominated reefs attached to hard substrata. Interestingly, we repeatedly recover *H. magnifica* and *S. gigantea* within the same group as well (Figures 2 and 3), but with little support. However, like *S. haddoni* and *S. mertensii*, *H. magnifica* and *S. gigantea* also occupy distinct habitats. *Stichodactyla gigantea* is described to occupy shallow sandy-bottom habitats (while *S. haddoni* is described to occur at greater depths), and *H. magnifica* occupies coral-dominated reef habitat attached directly to hard substrata similarly to *S. mertensii*. Like the Heteractina above, targeted phylogenetic analyses of Stichodactylina using highly resolving genomic markers should accompany morphological revisions to resolve hierarchical relationships within the clade, and explore whether there have been repeated ecological speciation events within the group.

Although our data show *E. quadricolor* to be monophyletic, the branch lengths and topological support within our focused reconstruction of this species (Figure 5) suggest that this grouping may be a species complex. The ((Red Sea, Maldives), Tonga) topological relationship is a classic Indian Ocean/Pacific Ocean biogeographic pattern found at both the intra-and interspecific levels in the phylogeographic literature (reviewed by Bowen et al., 2016). However, co-occurring samples from Japan also belong to both of these well-supported clades (Figure 5), and provide further evidence these may represent cryptic species. The long branch linking samples from the United Arab Emirates with the rest of the Indo-Pacific can also likely be construed as a third cryptic lineage. Counterintuitively, the ability of the current suite of genetic markers we used in this study to resolve hierarchical relationships within *E. quadricolor* may provide the strongest support for the distinctiveness of these lineages. Anthozoan mtDNA is largely uninformative at the intraspecific-level due to its slow rate of evolution (e.g. Daly et al, 2010; Shearer et al., 2002), and the combination of mtDNA and nuDNA markers used here are regularly unable to differentiate between species within the same genus (e.g. Daly et al., 2017; Grajales & Rodríguez, 2016; Larson & Daly, 2016; Rodríguez et al., 2014). However, sympatric species within the genus *Exaiptasia* have been delimited previously using these markers (Grajales & Rodríguez, 2016), and taken together, we provide the first preliminary evidence that there may be more than 10 species of clownfish-hosting sea anemones. As *E. quadricolor* hosts the greatest number of clownfish species throughout the Indo-Pacific (Fautin & Allen, 1992), the presence of extensive undescribed cryptic anemone species within this group could have wide ranging implications for our understanding of clownfish host associations, their symbiotic specialist/generalist designations, and to the degree to which these mutualistic relationships are nested (e.g. Ollerton et al., 2007).

In summary, our data conclusively reject the current taxonomy of the clownfish-hosting sea anemones. Major revisionary work at the familial, generic, and specific level is required to align taxonomy of the clownfish-hosting anemones with the phylogenetic results we present here.

Although we reject the current taxonomy of the clownfish-hosting sea anemones, we fail to reject the hypothesis that symbiosis with clownfishes evolved multiple times within Actiniaria. While it could be argued that our analysis cannot distinguish between a single or multiple origins of symbiosis with clownfishes, our likelihood ratio test convincingly refutes a model with only a single origin. Thus, we interpret our phylogenetic reconstruction to show that symbiosis with clownfishes evolved, minimally, three times within Actiniaria— all within the superfamily Actinioidea. We identify these independent origins as 1) within Stichodactylina, after *S. helianthus* diverged in the Tropical Western Atlantic, 2) in Heteractina, and 3) in the *E. quadricolor* species complex. We identify *Thalassianthus* as a possible lineage in which symbiosis with clownfishes was lost. This interpretation is made with the important caveat that clownfishes would have had to have been symbiotic with the common ancestor of these clades, with this relationship being retained as these lineages further diversified. If diversification of all 10 nominal anemone hosts pre-dates the onset of their symbiosis with fishes, then we would interpret the symbiosis to have evolved independently within each host species (i.e. 10 independent evolutionary origins).

The lack of a fossil record for anemones broadly and unknown mutation rates for the loci we have used here prevents us from producing time-calibrated phylogenetic trees. However, biogeographic patterns can help provide a minimum age for some of our clades. The phylogenetic position of the non-hosting, Tropical Western Atlantic species *S. helianthus* within Stichodactylina suggests that this clade has a Tethyan biogeographic origin, as it is divided into Atlantic and Indo-Pacific lineages (Figures 2 & 3). The final closure of the Tethys Sea, termed the Terminal Tethyan Event (TTE), occurred during the Miocene and has been dated to between 12-18 myr (reviewed by Cowman & Bellwood, 2013). Although vicariance between Atlantic and Indo-Pacific lineages has been shown to include numerous pulses of diversification that often pre-date the TTE in other taxonomic groups (Cowman & Bellwood, 2013), this event establishes a minimum potential age on the Stichodactylina and for the adoption of the symbiosis, which is inferred to have occurred at or after the split between *S. helianthus* and the rest of Stichodactylina. Further, within *E. quadricolor*, because anemones from the United Arab Emirates are interpreted as sister to the rest of the Indo-Pacific, we also infer a Tethyan origin and minimum age of 12-18 myr for *E. quadricolor*. Tethyan relic species found around the Arabian Peninsula, such as cowries and opisthobranch molluscs, show similar phylogenetic structure (e.g. Malaquias & Reid, 2009; Meyer, 2003). Interestingly, the root of the clownfish clade is consistently dated between 12-19 myr (e.g. Litsios et al., 2012; Litsios et al., 2014a; Rolland et al., 2018), which broadly coincides with the minimum potential age of the Stichodactylina, the *E. quadricolor* clade and the TTE. The clownfish subfamily, however, shows strong support for a Coral Triangle biogeographic origin, followed by an independent geographical radiation in the Western Indian Ocean ~4 mya (Litsios et al., 2014a). Taken together, it appears probable that the biogeographic setting of diversification differed between clownfish and at least two major lineages of host anemones.

Like many obligate mutualisms, the clownfish-sea anemone symbiosis is often described in a manner in which it is presumed that the interacting species have co-evolved to varying degrees. Throughout their ranges, host anemones are rarely unoccupied, and indeed, anemones do not serve as mere habitat space for clownfish. Rather, they receive tangible benefits from their symbiotic partners that allow them to compete and maintain habitat space, ward off predators, and obtain a steady source of nitrogen in an oligotrophic environment (Cleveland et al., 2011; Dunn, 1981; Fautin, 1991; Roopin et al., 2008; Szczebak et al., 2013). The ecological success of the anemone hosts, particularly on low-latitude reefs, clearly relies on clownfish. Whether the diversity and evolutionary history of the host anemones has been similarly driven by their symbiosis with clownfish is less clear. Our ability to discern a Tethyan origin for the Stichodactylina, and possibly *E. quadricolor*, demonstrates that host anemones likely had to expand their geographic ranges from the Tethys Sea across the Indian Ocean and into the Coral Triangle as solitary individuals before establishing symbiosis with clownfishes. Further, if host anemone diversification into Atlantic and Indo-Pacific Ocean basins preceded the TTE, as may be the case in other groups with Tethyan biogeographic histories, it would imply that the majority of host anemone diversity arose before the clownfish radiation. Whether diversification of the clownfishes and host anemones is fully decoupled will require extensive biogeographic sampling across all host anemones. This appears warranted as *E. quadricolor* (and possibly *H*. *magnifica* and others) may be a species complex and host anemones could be far more diverse than we currently recognize. Beyond spatial patterns of diversification, our understanding of actiniarian diversity broadly would stand to benefit from calibrated genomic mutation rates, allowing us to explore temporal patterns of co-diversification as well. These will be valuable avenues for future studies employing highly resolving genomic markers coupled with increased biogeographic sampling for each nominal anemone host lineage.

## Acknowledgements

We thank the Small Island Research Station (Fares-Maathoda, Maldives) for field research support and logistics, especially Mohamed Aslam, Ali Zahir, and Rahula Suhail. Kevin Kohen (Live Aquaria) and Laura Simmons (Cairns Marine) provided anemone samples from Tonga and Australia. Lily Berniker (AMNH) helped with sample handling and accession. Samples from the Philippines were collected with field support from Terry Gosliner, Rich Mooi, and Chrissy Piotrowski (California Academy of Science).

## Ethics

Sea anemones and tissue samples were collected under research permits: 30-D/INDIV/2018/27 (Maldives) and RE-17-04 and RE-18-07 (Palau). From the United Arab Emirates, tissue samples were collected and exported with official written permission from M. Fathima Al Antubi, head of the Environmental Protection Department, Government of Fujairah, Dibba Municipality. Samples from the Philippines were collected with the support of the Philippines Department of Agriculture, Bureau of Fisheries and Aquatic Resources, and the National Fisheries Research and Development Institute of the Philippines under permits GP-0085-15 and GP-0077-14, and are loaned to MD via Materials Transfer Agreement between her and the California Academy of Sciences. No permits were required to collect sea anemones in Japan.

## Data, code, and materials

All data are available in the Dryad Digital Repository: doi:XXXXX

## Author contributions

BMT conceived the project, collected tissue samples and field data, performed laboratory benchwork and sequencing, performed analyses, and wrote the paper. CB, RL, LCG, VVD, and TC performed laboratory bench-work and sequencing, performed analyses, and critically revised the paper. CPM, MLB, AB, KY, JDR, TF, MD collected tissue samples and field data, and critically revised the paper. ER conceived the project and critically revised the paper.

