## Supplementary Material for "Phylogenetic relationships among the clownfish-hosting sea anemones"

**Table S1.** List of non-clownfish hosting sea anemones and GenBank Accession numbers used in this study. Individuals are listed alphabetically by Superfamily, Family, Genus, and Species.

| **Superfamily** | **Family** | ***Genus*** | ***Species*** | **Cox3** | **12S** | **16S** | **18S** | **28S** |
| --- | --- | --- | --- | --- | --- | --- | --- | --- |
| Actinernoidea | Actinernidae | *Actinernus* | *antarcticus* | ------------ | KJ482930 | KJ482966 | KJ483023 | KJ483126 |
| Actinernoidea | Actinernidae | *Isactinernus* | *quadrolobatus* | KJ482998 | KJ482932 | KJ482968 | KJ483024 | KJ483105 |
| Actinernoidea | Actinernidae | *Synhalcurias* | *elagans* | ------------- | KJ482942 | ------------ | KJ483021 | KJ483120 |
| Actinernoidea | Halcuriidae | *Halcurias* | *pilatus* | KJ482997 | KJ482931 | KJ482967 | KJ483020 | KJ483109 |
| Actinioidea | Actiniidae | *Actinia* | *fragacea* | GU473334 | EU190714 | EU190756 | EU190845 | KJ483085 |
| Actinioidea | Actiniidae | *Actinia* | *tenebrosa* | KT852330 | KT852045 | KT852111 | KT852174 | ------------ |
| Actinioidea | Actiniidae | *Anemonia* | *erythraea* | KY789271 | KY789302 | KY789335 | ------------ | ------------ |
| Actinioidea | Actiniidae | *Anemonia* | *viridis* | GU473335 | EU190718 | EU190760 | EU190849 | KJ483095 |
| Actinioidea | Actiniidae | *Anthopleura* | *annae* | KY789293 | KY789327 | KY789360 | ------------ | KY789392 |
| Actinioidea | Actiniidae | *Anthopleura* | *artemisia* | KT852300 | KT852015 | KT852081 | KT852148 | ------------ |
| Actinioidea | Actiniidae | *Anthopleura* | *atodai* | KT852275 | KT851993 | KT852055 | KT852123 | KT852247 |
| Actinioidea | Actiniidae | *Anthopleura* | *ballii* | KY789281 | KY789311 | KY789346 | ------------ | KY789376 |
| Actinioidea | Actiniidae | *Anthopleura* | *biscayensis* | KY789284 | KY789315 | KY789350 | ------------ | KY789380 |
| Actinioidea | Actiniidae | *Anthopleura* | *buddemeieri* | ------------ | KY789316 | KY789351 | ------------ | KY789381 |
| Actinioidea | Actiniidae | *Anthopleura* | *dixoniana* | KY789276 | KY789307 | KY789341 | ------------ | ------------ |
| Actinioidea | Actiniidae | *Anthopleura* | *dowii* | KY789286 | KY789318 | KY789353 | ------------ | KY789383 |
| Actinioidea | Actiniidae | *Anthopleura* | *elegantissima* | GU473333 | EU190713 | EU190755 | EU190844 | KT852248 |
| Actinioidea | Actiniidae | *Anthopleura* | *fuscoviridis* | KY789272 | KY789303 | KY789336 | ------------ | KY789369 |
| Actinioidea | Actiniidae | *Anthopleura* | *handi* | KT852298 | KT852013 | KT852079 | KT852146 | KY789387 |
| Actinioidea | Actiniidae | *Anthopleura* | *insignis* | KY789297 | KY789331 | KY789364 | ------------ | KY789395 |
| Actinioidea | Actiniidae | *Anthopleura* | *krebsi* | KY789275 | KY789305 | KY789339 | ------------ | KY789372 |
| Actinioidea | Actiniidae | *Anthopleura* | *kurogane* Korea | KY789288 | KY789321 | KY789355 | ------------ | KY789385 |
| Actinioidea | Actiniidae | *Anthopleura* | *midori* | KY789289 | KY789324 | ------------ | ------------ | KY789388 |
| Actinioidea | Actiniidae | *Anthopleura* | *nigrescen* | KY789278 | KY789306 | KY789343 | ------------ | KY789373 |
| Actinioidea | Actiniidae | *Anthopleura* | *rosea* | KT852324 | KT852039 | KT852104 | KT852168 | ------------ |
| Actinioidea | Actiniidae | *Anthopleura* | sp*. "inornata"* | KY789274 | KY789304 | KY789338 | ------------ | KY789371 |
| Actinioidea | Actiniidae | *Anthopleura* | *thallia* | KY789300 | KY789333 | KY789366 | ------------ | KY789397 |
| Actinioidea | Actiniidae | *Anthopleura* | *waridi* | KY789270 | KY789301 | KY789334 | ------------ | KY789368 |
| Actinioidea | Actiniidae | *Anthostella* | *stephensoni* | JQ810726 | JQ810719 | JQ810721 | JQ810723 | KJ483132 |
| Actinioidea | Actiniidae | *Aulactinia* | *incubans* | KT852299 | KT852014 | KT852080 | KT852147 | KT852256 |
| Actinioidea | Actiniidae | *Aulactinia* | *marplatensis* | KT852281 | KT851999 | KT852061 | KT852129 | KT852249 |
| Actinioidea | Actiniidae | *Aulactinia* | *stella* | KT852329 | KT852044 | KT852110 | KT852173 | KT852263 |
| Actinioidea | Actiniidae | *Aulactinia* | *vancouverensis* | KT852305 | KT852019 | KT852085 | KT852151 | ------------ |
| Actinioidea | Actiniidae | *Aulactinia* | *veratra* | KT852283 | KT852001 | KT852063 | KT852131 | ------------ |
| Actinioidea | Actiniidae | *Bolocera* | *kerguelensis* | KJ482985 | KJ482925 | KJ482965 | KJ483029 | KJ483133 |
| Actinioidea | Actiniidae | *Bunodactis* | *reynaudi* | KT852326 | KT852041 | KT852106 | KT852170 | KT852260 |
| Actinioidea | Actiniidae | *Bunodactis* | *verrucosa* | FJ489484 | EU190723 | EU190766 | EU190854 | EU190812 |
| Actinioidea | Actiniidae | *Bunodosoma* | *cavernatum* | KY789282 | KY789313 | KY789348 | ------------ | KY789378 |
| Actinioidea | Actiniidae | *Bunodosoma* | *grandis* | GU473336 | EU190722 | EU190765 | EU190853 | EU190811 |
| Actinioidea | Actiniidae | *Bunodosoma* | *granuliferum* | KY789283 | KY789314 | KY789349 | ------------ | KY789379 |
| Actinioidea | Actiniidae | *Epiactis* | *australiensis* | KT852282 | KT852000 | KT852062 | KT852130 | ------------ |
| Actinioidea | Actiniidae | *Epiactis* | *fernaldi* | KT852288 | KT852005 | KT852068 | KT852136 | KT852252 |
| Actinioidea | Actiniidae | *Epiactis* | *georgiana* | KT852290 | KT852007 | KT852070 | KT852138 | KT852254 |
| Actinioidea | Actiniidae | *Epiactis* | *handi* | KT852269 | KT851988 | KT852050 | KT852118 | KT852245 |
| Actinioidea | Actiniidae | *Epiactis* | *handi* | KT852271 | KT851990 | KT852052 | KT852120 | KT852268 |
| Actinioidea | Actiniidae | *Epiactis* | *japonica* | KT852272 | KT851991 | KT852053 | KT852121 | ------------ |
| Actinioidea | Actiniidae | *Epiactis* | *japonica* | KT852273 | KT851992 | KT852054 | KT852122 | ------------ |
| Actinioidea | Actiniidae | *Epiactis* | *japonica* | KY789285 | KY789317 | KY789352 | ------------ | KY789382 |
| Actinioidea | Actiniidae | *Epiactis* | *lisbethae* | KT852289 | KT852006 | KT852069 | KT852137 | KT852253 |
| Actinioidea | Actiniidae | *Epiactis* | *lisbethae* | GU473360 | EU190727 | EU190771 | EU190858 | EU190816 |
| Actinioidea | Actiniidae | *Epiactis* | *prolifera* | KT852270 | KT851989 | KT852051 | KT852119 | KT852246 |
| Actinioidea | Actiniidae | *Epiactis* | *ritteri* | KT852276 | KT851994 | KT852056 | KT852124 | ------------ |
| Actinioidea | Actiniidae | *Epiactis* | *ritteri* | KT852277 | KT851995 | KT852057 | KT852125 | ------------ |
| Actinioidea | Actiniidae | *Epiactis* | *thompsoni* | KT852293 | KT852010 | KT852073 | KT852141 | ------------ |
| Actinioidea | Actiniidae | *Glyphoperidium* | *bursa* | KJ482982 | KJ482923 | KJ482961 | KJ483033 | KJ483136 |
| Actinioidea | Actiniidae | *Gyractis* | *sesere* | KT852297 | KT852012 | KT852078 | KT852145 | KY789386 |
| Actinioidea | Actiniidae | *Isactinia* | *olivacea* | KT852296 | ------------ | KT852077 | KT852144 | ------------ |
| Actinioidea | Actiniidae | *Isosicyonis* | *alba* | KJ482981 | ------------ | KJ482959 | KJ483030 | KJ483134 |
| Actinioidea | Actiniidae | *Isosicyonis* | *striata* | FJ489493 | EU190736 | EU190781 | EU190864 | FJ489463 |
| Actinioidea | Actiniidae | *Isotealia* | *antarctica* | JQ810727 | JQ810720 | JQ810722 | ------------ | ------------ |
| Actinioidea | Actiniidae | *Korsaranthus* | *natalinesis* | KJ482987 | KJ482920 | KJ482958 | KJ483017 | KJ483117 |
| Actinioidea | Actiniidae | *Oulactis* | *muscosa* | KT852317 | KT852033 | KT852097 | KT852162 | KY789391 |
| Actinioidea | Actiniidae | *Phlyctenactis* | *tuberculosa* | KY789292 | KY789326 | KY789359 | ------------ | ------------ |
| Actinioidea | Actiniidae | *Pseudactinia* | *varia* | KY789294 | KY789328 | KY789361 | ------------ | ------------ |
| Actinioidea | Actiniidae | *Urticina* | *coriacea* | GU473351 | GU473282 | EU190797 | EU190877 | EU190840 |
| Actinioidea | Actiniidae | *Urticina* | *crassicornis* | KT852279 | KT851997 | KT852059 | KT852127 | ------------ |
| Actinioidea | Actiniidae | *Urticina* | *fecunda* | KT852287 | KT852004 | KT852067 | KT852135 | ------------ |
| Actinioidea | Actiniidae | *Urticina* | *grebelnyli* | KT852318 | KT852034 | KT852098 | KT852163 | ------------ |
| Actinioidea | Actinodendridae | *Actinostephanus* | *haeckeli* | GU473353 | KJ482936 | EU190762 | KJ483034 | EU190808 |
| Actinioidea | Capneidae | *Capnea* | *georgiana* | KJ482990 | ------------ | KJ482951 | KJ483022 | KJ483050 |
| Actinioidea | Condylanthidae | *Charisea* | *saxicola* | KT852306 | KT852020 | KT852086 | KT852152 | ------------ |
| Actinioidea | Haloclavidae | *Haloclava* | sp*.* | KJ482989 | KJ482924 | KJ482963 | KJ483031 | KJ483138 |
| Actinioidea | Haloclavidae | *Haloclava* | *producta* | JF833008 | EU190734 | EU190779 | AF254370 | EU190823 |
| Actinioidea | Haloclavidae | *Harenactis* | *argentina* | KJ482984 | KJ482926 | KJ482964 | KJ483026 | KJ483047 |
| Actinioidea | Haloclavidae | *Peachia* | *cylindrica* | ------------ | EU190743 | EU190789 | KJ483015 | EU190732 |
| Actinioidea | Haloclavidae | *Stephanthus* | *antarcticus* | KJ482983 | KJ482927 | KJ482960 | KJ483019 | KJ483092 |
| Actinioidea | Liponematidae | *Liponema* | *brevicornis* | KJ483001 | EU190738 | EU190784 | EU190866 | KJ483139 |
| Actinioidea | Liponematidae | *Liponema* | *multiporum* | ------------ | KJ482922 | KJ482962 | ------------ | ------------ |
| Actinioidea | Phymanthidae | *Phymanthus* | *loligo* | GU473345 | EU190745 | EU190791 | EU190871 | ------------ |
| Actinioidea | Phymanthidae | *Phymanthus* | *crucifer* | KJ910346 | KJ910343 | KJ910345 | MH670399 | MH670928 |
| Actinioidea | Phymanthidae |  |  | KJ910347 | KJ910343 | KJ910345 | MH670400 | MH670930 |
| Actinioidea | Phymanthidae |  |  | KJ910348 | KJ910343 | KJ910345 | MH670401 | MH670935 |
| Actinioidea | Preactiidae | *Preactis* | *milliardae* | KJ482986 | KJ482921 | KJ482957 | KJ483018 | KJ483118 |
| Actinioidea | Preactiidae | *Dactylanthus* | *antarcticus* | GU473358 | GU473272 | AY345877 | AF052896 | KJ483086 |
| Actinioidea | Thalassianthidae | *Thalassianthus* | *aster* | KC812240 | ------------ | ------------ | KC812195 | KC812219 |
| Actinioidea | Thalassianthidae | *Thalassianthus* | *aster* | KC812241 | KC812146 | KC812167 | KC812196 | KC812220 |
| Actinioidea | Thalassianthidae | *Thalassianthus* | *hemprichii* | KC812238 | ------------ | KC812168 | KC812193 | KC812217 |
| Actinioidea | Thalassianthidae | *Thalassianthus* | *hemprichii* | KC812239 | KC812145 | KC812169 | KC812194 | KC812218 |
| Actinostoloidea | Actinostolidae | *Actinostola* | *chilensis* | GU473357 | ------------ | GU473285 | GU473302 | KJ483110 |
| Actinostoloidea | Actinostolidae | *Actinostola* | *crassicornis* | GU473332 | ------------ | EU190753 | EU190843 | EU272904 |
| Actinostoloidea | Actinostolidae | *Actinostola* | *georgiana* | KJ482991 | KJ482928 | KJ482952 | KJ483032 | KJ483099 |
| Actinostoloidea | Actinostolidae | *Antholoba* | *achates* | GU473356 | GU473269 | GU473284 | GU473301 | GU473318 |
| Actinostoloidea | Actinostolidae | *Anthosactis* | *janmayeni* | GU473363 | KJ482938 | GU473292 | GU473308 | GU473324 |
| Actinostoloidea | Actinostolidae | *Hormosoma* | *scotti* | GU473366 | EU190733 | EU190778 | EU190863 | EU190822 |
| Actinostoloidea | Actinostolidae | *Paranthus* | *niveus* | GU473344 | GU473277 | GU473295 | GU473311 | GU473327 |
| Actinostoloidea | Actinostolidae | *Stomphia* | *didemon* | GU473348 | EU190749 | EU190795 | EU190875 | EU190837 |
| Actinostoloidea | Actinostolidae | *Stomphia* | *selaginella* | GU473349 | GU473280 | GU473298 | GU473314 | GU473331 |
| Edwardsioidea | Edwardsiidae | *Edwardsia* | *elegans* | GU473338 | EU190726 | EU190770 | EU190857 | EU190815 |
| Edwardsioidea | Edwardsiidae | *Edwardsia* | *japonica* | GU473359 | GU473274 | GU473288 | GU473304 | KJ483048 |
| Edwardsioidea | Edwardsiidae | *Edwardisella* | *loveni* | KX946217.1 | KX946216.1 | KX946212 | KX946218 | KX946219 |
| Edwardsioidea | Edwardsiidae | *Edwardsia* | *timida* | KJ482996 | GU473281 | KT852113 | GU473315 | KJ483088 |
| Edwardsioidea | Edwardsiidae | *Edwardsianthus* | *gilbertensis* | ------------- | EU190728 | EU190772 | EU190859 | EU190817 |
| Edwardsioidea | Edwardsiidae | *Nematostella* | *vectensis* | FJ489501 | EU190750 | AY169370 | AF254382 | KJ483089 |
| Metridioidea | Actinoscyphiidae | *Actinoscyphia* | *plebeia* | FJ489476 | EU190712 | EU190754 | FJ489437 | EU190800 |
| Metridioidea | Aiptasiidae | *Aiptasia* | *couchii* | KP761405 | KP761199 | KP761254 | KP761301 | ------------ |
| Metridioidea | Aiptasiidae | *Aiptasia* | *couchii* | KP761403 | KP761200 | KP761255 | KP761303 | ------------ |
| Metridioidea | Aiptasiidae | *Aiptasia* | *mutabilis* | FJ489505 | JF832963 | FJ489418 | FJ489438 | FJ489469 |
| Metridioidea | Aiptasiidae | *Aiptasia* | *mutabilis* | KP761404 | KP761194 | KP761248 | KP761300 | ------------ |
| Metridioidea | Aiptasiidae | *Aiptasia* | *pulchella* | FJ489477 | EU190715 | EU190757 | EU190846 | EU190803 |
| Metridioidea | Aiptasiidae | *Aiptasiogeton* | *hyalinus* | ------------ | KR704266 | KR186040 | KR704268 | ------------ |
| Metridioidea | Aiptasiidae | *Bartholomea* | *annulata* | FJ489483 | EU190721 | EU190763 | EU190851 | KJ483068 |
| Metridioidea | Aiptasiidae | *Bellactis* | *ilkalyseae* | KP761393 | KR186020 | KR186036 | KR186051 | ------------ |
| Metridioidea | Aiptasiidae | *Bellactis* | *ilkalyseae* | ------------ | KR186021 | KR186037 | KR186052 | ------------ |
| Metridioidea | Aiptasiidae | *Exaiptasia* | *brasiliensis* | KP761386 | KP761188 | KP761239 | KP761312 | ------------ |
| Metridioidea | Aiptasiidae | *Exaiptasia* | *pallida* | KP761323 | KP761184 | KP761283 | ------------ | ------------ |
| Metridioidea | Aiptasiidae | *Exaiptasia* | *pallida* | KP761361 | KP761183 | KP761270 | KP761286 | ------------ |
| Metridioidea | Aiptasiidae | *Exaiptasia* | *pallida* | ------------ | KP761182 | ------------ | KP761279 | ------------ |
| Metridioidea | Aiptasiidae | *Laviactis* | *lucida* | KP761402 | KP761192 | KP761243 | KP761296 | ------------ |
| Metridioidea | Aiptasiidae | *Neoaiptasia* | *morbilla* | JF833010 | EU190742 | EU190788 | EU190869 | EU190831 |
| Metridioidea | Aliciidae | *Alicia* | *mirabilis* | KP761410 | KP761213 | ------------ | KP761310 | KP761329 |
| Metridioidea | Aliciidae | *Alicia* | *sansibarensis* | KJ483000 | KJ482933 | KJ482953 | KJ483016 | KJ483116 |
| Metridioidea | Aliciidae | *Triactis* | *producta* | GU473350 | EU490525 | ------------ | EU190876 | EU190839 |
| Metridioidea | Amphianthidae | *Amphianthus* | sp. | FJ489502 | FJ489413 | FJ489432 | FJ489450 | FJ489467 |
| Metridioidea | Amphianthidae | *Peronanthus* | sp*.* | KJ482976 | KJ482917 | KJ482956 | KJ483014 | KJ483066 |
| Metridioidea | Andvakiidae | *Andvakia* | *boninensis* | FJ489479 | EU190717 | EU190759 | EU190848 | EU190805 |
| Metridioidea | Andvakiidae | *Andvakia* | *discipulorum* | ------------ | GU473273 | GU473287 | GU473316 | GU473320 |
| Metridioidea | Andvakiidae | *Telmatactis* | sp. | ------------ | JF832968 | JF832979 | KJ483013 | JF833001 |
| Metridioidea | Antipodactinidae | *Antipodactis* | *awii* | GU473337 | GU473271 | GU473286 | GU473303 | GU473319 |
| Metridioidea | Bathyphelliidae | *Bathyphellia* | *australis* | FJ489482 | FJ489402 | FJ489422 | EF589063 | EF589086 |
| Metridioidea | Boloceroididae | *Bunodeopsis* | *globulifera* | KJ482992 | KJ482940 | KJ482949 | KJ483025 | KJ483122 |
| Metridioidea | Diadumenidae | *Diadumene* | *cincta* | FJ489490 | EU190725 | EU190769 | EU190856 | EU190814 |
| Metridioidea | Diadumenidae | *Diadumene* | *leucolena* | JF833006 | JF832957 | JF832977 | JF832986 | KJ483123 |
| Metridioidea | Diadumenidae | *Diadumene* | sp. | JF833005 | JF832960 | JF832976 | JF832980 | KJ483130 |
| Metridioidea | Diadumenidae | *Diadumene* | *lineata* | JF833007 | JF832965 | JF832973 | JF832987 | JF832998 |
| Metridioidea | Diadumenidae | *Diadumene* | *lineata* | FJ489506 | EU190730 | EU190774 | EU190860 | EU190819 |
| Metridioidea | Galantheanthemidae | *Galatheanthemum* | *profundus* | KJ482978 | KJ482919 | KJ482954 | KJ483011 | KJ483119 |
| Metridioidea | Galantheanthemidae | *Galatheanthemum* | sp | KJ482977 | KJ482918 | KJ482955 | KJ483012 | KJ483065 |
| Metridioidea | Halcampidae | *Cactosoma* | sp | GU473346 | GU473279 | GU473297 | GU473313 | GU473329 |
| Metridioidea | Halcampidae | *Halcampa* | *duodecimcirrata* | ------------ | JF832966 | EU190776 | AF254375 | EU190820 |
| Metridioidea | Halcampidae | *Halcampoides* | *purpureus* | ------------ | EU190735 | EU190780 | AF254380 | EU190824 |
| Metridioidea | Hormathiidae | *Actinauge* | *richardi* | FJ489480 | EU190719 | EU190761 | EU190850 | EU190807 |
| Metridioidea | Hormathiidae | *Adamsia* | *palliata* | FJ489474 | FJ489398 | FJ489419 | FJ489436 | FJ489452 |
| Metridioidea | Hormathiidae | *Allantactis* | *parasitica* | FJ489478 | FJ489399 | FJ489420 | FJ489439 | FJ489454 |
| Metridioidea | Hormathiidae | *Calliactis* | *japonica* | FJ489486 | FJ489403 | FJ489423 | FJ489441 | FJ489456 |
| Metridioidea | Hormathiidae | *Calliactis* | *parasitica* | FJ489475 | EU190711 | EU190752 | EU190842 | EU190799 |
| Metridioidea | Hormathiidae | *Calliactis* | *polypus* Hawaii | FJ489485 | FJ489407 | FJ489427 | FJ489445 | FJ489459 |
| Metridioidea | Hormathiidae | *Calliactis* | *tricolor* | FJ489488 | FJ489405 | FJ489425 | FJ489443 | FJ489458 |
| Metridioidea | Hormathiidae | *Chondrophellia* | *orangina* | FJ489489 | FJ489406 | FJ489426 | FJ489444 | KJ483060 |
| Metridioidea | Hormathiidae | *Cricophorus* | *nutrix* | KT852286 | ------------ | KT852066 | KT852134 | ------------ |
| Metridioidea | Hormathiidae | *Hormathia* | *armata* | FJ489491 | EU190731 | EU190775 | EU190861 | FJ489460 |
| Metridioidea | Hormathiidae | *Hormathia* | *lacunifera* | FJ489492 | FJ489409 | FJ489428 | FJ489446 | FJ489461 |
| Metridioidea | Hormathiidae | *Hormathia* | *pectinata* | FJ489497 | FJ489415 | FJ489430 | FJ489448 | FJ489465 |
| Metridioidea | Hormathiidae | *Paracalliactis* | *japonica* | FJ489496 | FJ489411 | FJ489429 | FJ489447 | FJ489464 |
| Metridioidea | Hormathiidae | *Paraphelliactis* | *pabista.* | FJ489498 | FJ489412 | FJ489431 | FJ489449 | FJ489466 |
| Metridioidea | Isanthidae | *Isanthus* | *capensis* | GU473362 | JF832967 | GU473291 | GU473291 | GU473323 |
| Metridioidea | Isanthidae | *Isoparactis* | *fabiani* | GU473355 | JF832964 | GU473283 | GU473300 | KJ483124 |
| Metridioidea | Isanthidae | *Isoparactis* | *fionae* | KC700007 | KC700001 | KC700003 | KC700004 | ------------ |
| Metridioidea | Isanthidae | *Isoparactis* | *ferax* | KC700008 | KC700002 | ------------ | KC700005 | KC700006 |
| Metridioidea | Kadosactinidae | *Alvinactis* | *chessi* | GU473352 | GU473278 | GU473296 | GU473312 | GU473328 |
| Metridioidea | Kadosactinidae | *Cyananthea* | *hourdezi* | GU473364 | GU473275 | GU473293 | GU473309 | GU473325 |
| Metridioidea | Kadosactinidae | *Jasonactis* | *erythraios* | GU473339 | ------------ | GU473289 | GU473305 | GU473330 |
| Metridioidea | Kadosactinidae | *Kadosactis* | *antarctica* | FJ489504 | FJ489410 | EU190782 | EU190865 | EU190825 |
| Metridioidea | Metridiidae | *Metridium* | *senile fimbriatum* | KT852309 | KT852023 | KT852089 | JF832988.1 | KT852257 |
| Metridioidea | Metridiidae | *Metridium* | *s. fibratum* (Japan) | JF833009 | ------------ | JF832974 | JF832988 | JF832996 |
| Metridioidea | Metridiidae | *Metridium* | *s. lobatum* (Argentina) | JF833002 | JF832962 | JF832971 | JF832981 | JF832991 |
| Metridioidea | Metridiidae | *Metridium* | *senile* | FJ489494 | EU190740 | EU190786 | AF052889 | EU190829 |
| Metridioidea | Metridiidae | *Metridium* | *senile* | KJ482975 | KJ482916 | KJ482950 | KJ483035 | KJ483113 |
| Metridioidea | Metridiidae | *Metridium* | *senile* | JF833003 | JF832961 | JF832972 | JF832982 | JF833000 |
| Metridioidea | Nemathidae | *Nemanthus* | *nitidus* | FJ489495 | EU190741 | EU190787 | EU190868 | EU190830 |
| Metridioidea | Ostiactinidae | *Ostiactis* | *pearseae* | GU473365 | EU190751 | EU190798 | EU190878 | EU190841 |
| Metridioidea | Phelliidae | *Phellia* | *exlex* | JF833004 | JF832958 | JF832978 | JF832984 | KJ483121 |
| Metridioidea | Phelliidae | *Phellia* | *gausapata* | FJ489473 | EU190744 | EU190790 | EU190870 | EU190833 |
| Metridioidea | Sagartiidae | *Actinothoe* | *sphyrodeta* | FJ489481 | FJ489401 | FJ489421 | FJ489440 | FJ489455 |
| Metridioidea | Sagartiidae | *Anthothoe* | *chilensis* | FJ489470 | FJ489397 | FJ489416 | FJ489434 | FJ489453 |
| Metridioidea | Sagartiidae | *Cereus* | *herpetodes* | ------------ | JF832956 | JF832969 | JF832983 | JF832992 |
| Metridioidea | Sagartiidae | *Cereus* | *pedunculatus* | FJ489471 | EU190724 | EU190767 | EU190855 | EU190813 |
| Metridioidea | Sagartiidae | *Sagartia* | *elegans* | JF833012 | ------------ | JF832970 | JF832989 | JF832994 |
| Metridioidea | Sagartiidae | *Sagartia* | *ornata* | JF833011 | JF832959 | JF832975 | JF832985 | JF832997 |
| Metridioidea | Sagartiidae | *Sagartia* | *troglodytes* | FJ489499 | EU190746 | KT852107 | EU190872 | KT852261 |
| Metridioidea | Sagartiidae | *Sagartiogeton* | *laceratus* | FJ489500 | EU190748 | EU190794 | EU190874 | KT852261 |
| Metridioidea | Sagartiidae | *Sagartiogeton* | *undatus* | FJ489472 | FJ489400 | FJ489417 | FJ489435 | FJ489462 |
| Metridioidea | Sagartiidae | *Verrillactis* | *paguri* | FJ489503 | FJ489414 | FJ489433 | FJ489440 | FJ489468 |
| **OUTGROUPS** |  |  |  |  |  |  |  |  |
| Corallimorpharia | Ricordeidae | *Ricordea* | *florida* | ------------ | KJ482913 | EF589057 | EF589067 | KJ483045 |
| Zoantharia | Parazoanthidae | *Parazoanthus* | *axinellae* | ----------- | GQ464940 | EU828754 | KC218416 | KJ483044 |

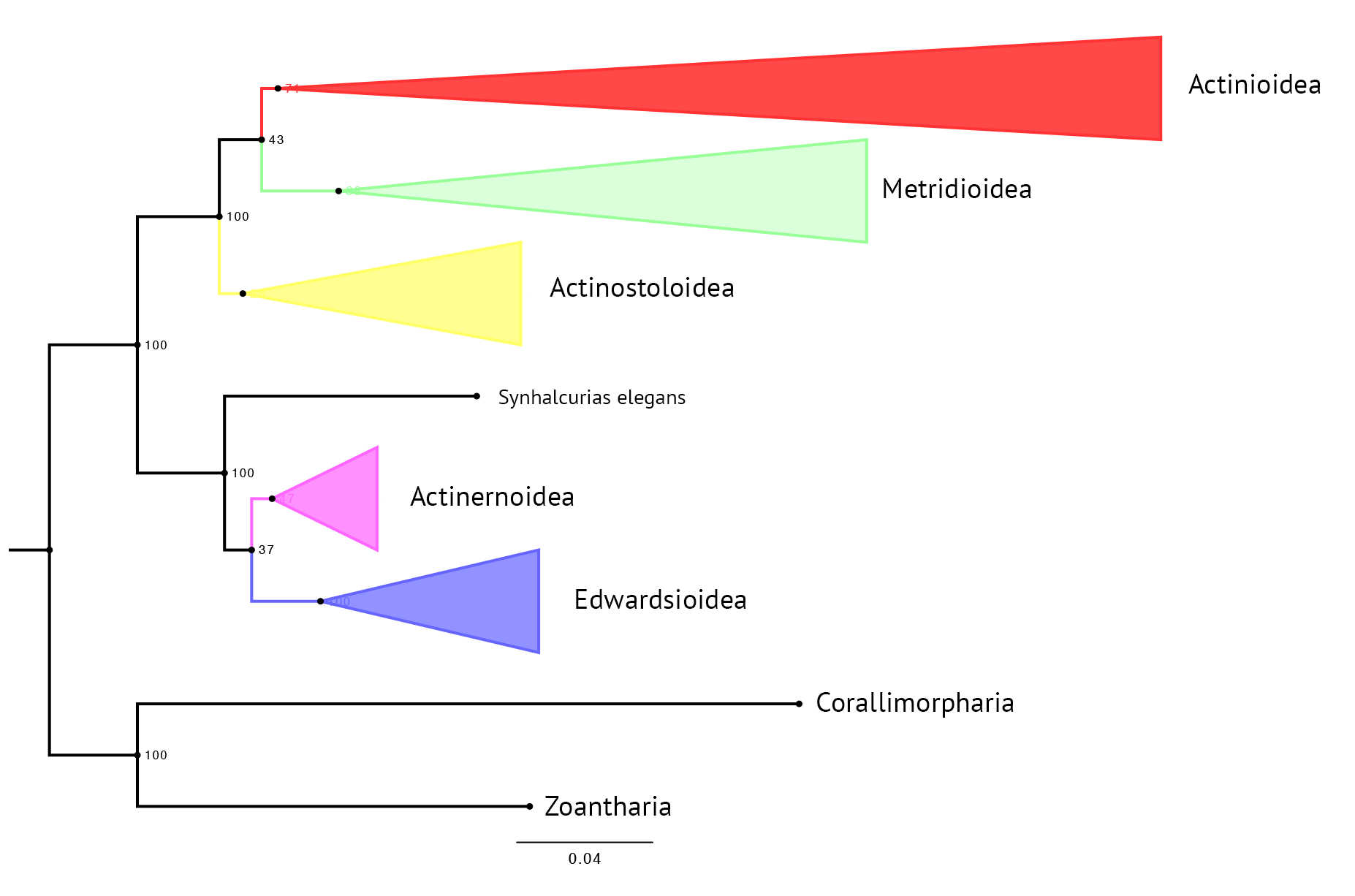

**Figure S1.** Maximum Likelihood (ML) phylogenetic reconstruction using the Actiniaria-wide dataset in RAxML. Tree reflects hierarchical relationships recovered among the anemone superfamilies (colored triangles). Size and shape of colored triangles reflects branch lengths within each superfamily. Numbers at each node represent bootstrap values from ML analyses.

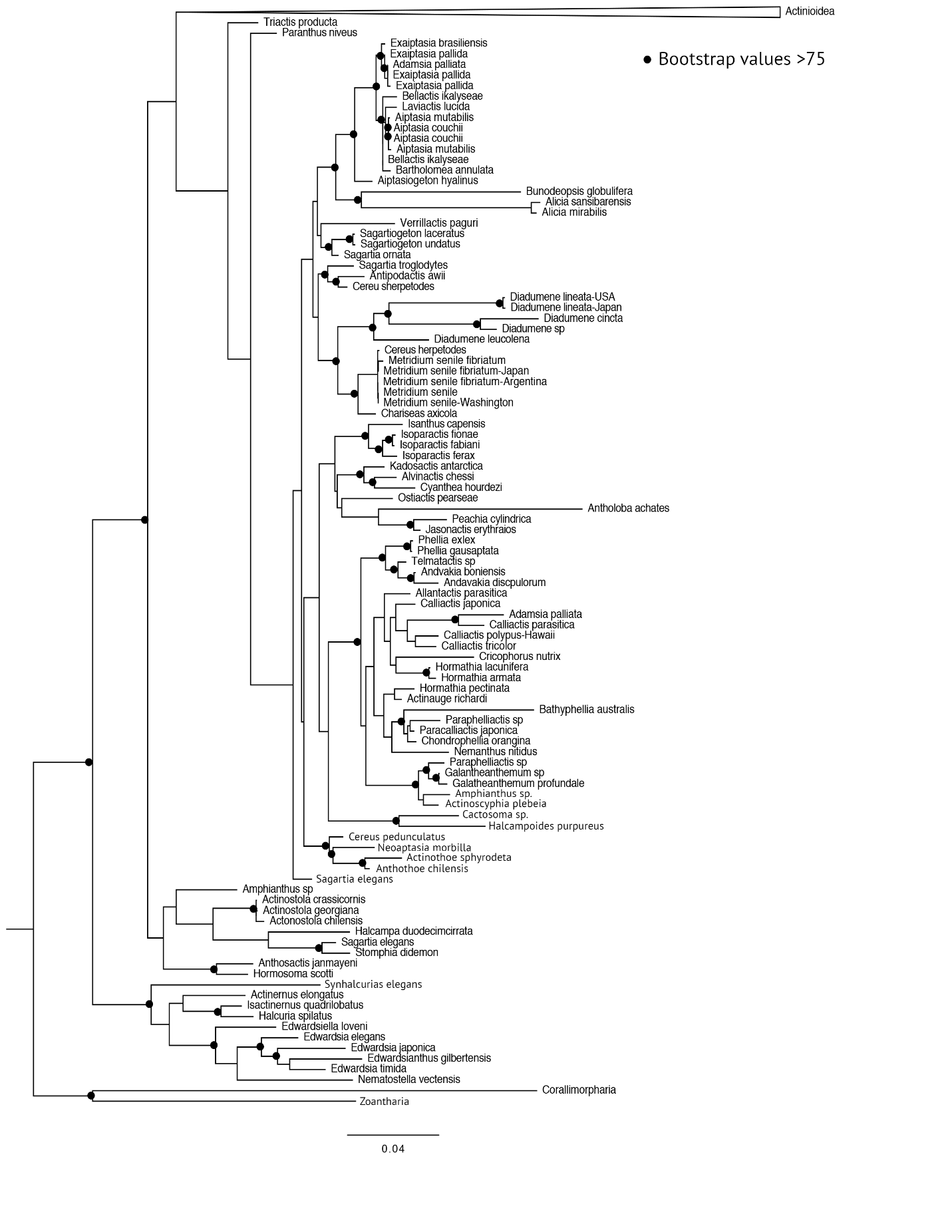

**Figure S2.** Maximum Likelihood (ML) phylogenetic reconstruction using the Actiniaria-wide dataset in RAxML. Tree reflects hierarchical relationships recovered among the anemone superfamilies Actinernoidea, Actinostoloidea, Edwardsioidea, and Metridioidea. The superfamily Actinioidea is collapsed for clarity; refer to Figure S3 for relationships within that clade. Black filled circles at nodes indicate bootstrap resampling values ≥75.

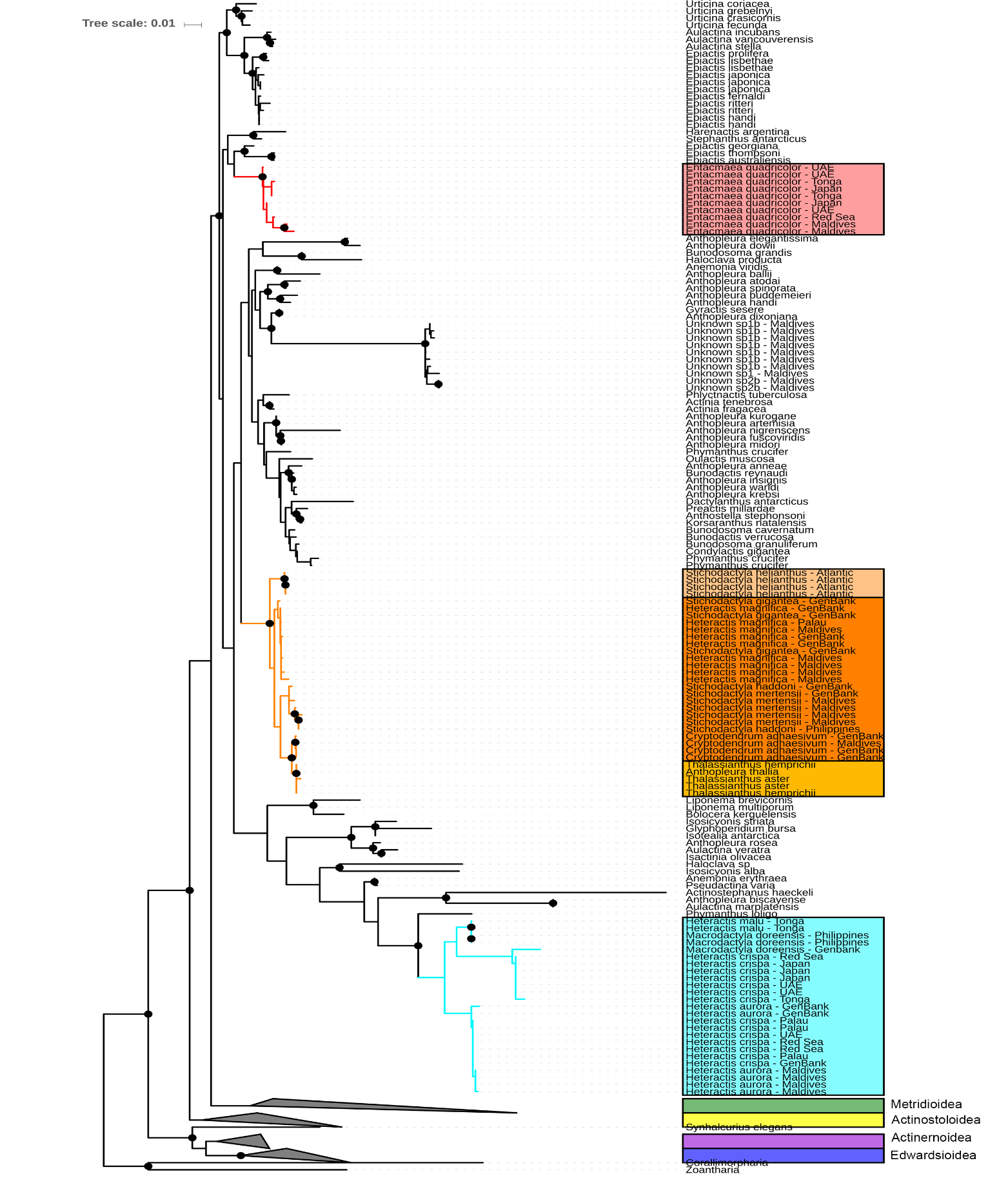

**Figure S3.** Maximum Likelihood (ML) phylogenetic reconstruction using the Actiniaria-wide dataset in RAxML. Tree reflects hierarchical relationships recovered among the anemone superfamily Actinioidea. Superfamilies Actinernoidea (purple), Actinostoloidea (yellow), Edwardsioidea (blue), and Metridioidea (green) collapsed for clarity. The figure highlights the three clades where symbiosis with clownfishes has evolved: Red = *Entacmaea quadricolor*, Light Blue = Heteractina (*Heteractis aurora*, *H. crispa*, *H. malu*, and *Macrodactyla doreensis*), Oranges = Stichodactylina (*Cryptodendrum adhaesivum, H. magnifica,* *Stichodactyla gigantea, S. haddoni, S. helianthus, S. mertensii, Thalassianthus aster, T. hemprichii*). Within Stichodactylina different hues of orange represent where *S. helianthus* diverged into the Atlantic Ocean, where symbiosis with clownfishes arose in the Indo-West Pacific, and where symbiosis with clownfishes may have been lost in the genus *Thalassianthus*. Black filled circles at nodes indicate bootstrap resampling values ≥ 75.
